# Fast receptor-antigen dissociation kinetics promote CAR-T cell tumor infiltration, functional fitness, and antitumor efficacy in solid tumors

**DOI:** 10.64898/2026.10.10.756030

**Authors:** Rui Mao, Huidong Shi, Mohamed S. Hussein, Bela Karia, Obed Koomson, Ramses Sadek, Vamsi Kota, Nicholas R.J. Gascoigne, Yukai He

## Abstract

**Background:** CAR-T cells have shown limited efficacy against solid tumors, where they face two major challenges: restricted tumor infiltration and sustained intensive engagement that drive them to exhaustion. Conventional CAR design has largely emphasized high-affinity interactions with slow dissociation, further limiting tumor infiltration and exacerbating exhaustion and dysfunction. We hypothesize that most current CARs are kinetically over-stabilized for solid tumors and that a fast-on/fast-off, “**cheek-kiss**” mode of engagement permits transient CAR–antigen disengagement, thereby facilitating tumor infiltration and preserving CAR-T cell fitness in tumor mass.

**Methods:** We identified two monoclonal antibodies targeting the same epitope of epidermal growth factor receptor with comparable *k*on but distinct *k*off rates: cetuximab (Cet) and its humanized derivative H1. CAR-T cells incorporating these binders were evaluated *in vitro* and *in vivo*. Large, established human tumor xenografts were treated with low-dose CAR-T cells in NSG mice to model conditions of limited T cell availability and sustained tumor antigen exposure.

**Results:** Cet and H1 exhibited comparable *k*on but markedly different *k*off rates, with H1 displaying fast-off kinetics with approximately 18-fold shorter binding dwell time. As CARs, H1 showed reduced antigen occupancy and less stable tumor-T cell engagement than Cet. Despite comparable *in vitro* cytotoxicity, H1 CAR-T cells accumulated more efficiently following tumor stimulation. At high doses, both CAR-T cell products eradicated established tumors. Under low-dose conditions, however, H1 CAR-T cells exhibited superior expansion, persistence, tumor infiltration, and antitumor efficacy, whereas Cet CAR-T cells showed limited expansion and only transient tumor control. Cet CAR-T cells progressively downregulated surface CAR expression, increased PD-1 expression, and exhibited reduced tumor infiltration and impaired functional recovery. In contrast, H1 CAR-T cells preserved surface CAR expression, exhibited greater tumor infiltration, maintained a less exhausted, more central memory-like phenotype, and retained effector function.

**Conclusion:** Fast-on/fast-off binding kinetics enhance CAR-T tumor infiltration, expansion, persistence, and functional fitness, resulting in improved control of established solid tumors. The data provides a rationale for shifting CAR design from affinity optimization toward kinetic optimization to improve the efficacy and safety of CAR-T therapy for solid tumors.

**WHAT IS ALREADY KNOWN ON THIS TOPIC:** CAR design has largely focused on antigen-binding affinity, whereas the contribution of binding kinetics, particularly the dissociation rate, to CAR-T cell function, expansion, persistence, tumor infiltration, and antitumor efficacy remains poorly defined.

**WHAT THIS STUDY ADDS:** In this study, we found that, compared with slow-off binding, fast-off kinetics enhance CAR-T cell tumor infiltration, expansion, persistence, and functional fitness in tumor-bearing mice, revealing previously underappreciated effects of antigen-binding kinetics on CAR-T cell tumor infiltration and performance in solid tumors.

**HOW THIS STUDY MIGHT AFFECT RESEARCH, PRACTICE OR POLICY:** This study provides a rationale for incorporating antigen-binding kinetics into CAR design and optimization. The TCR-inspired fast-on/fast-off antigen engagement may help CAR T cells maintain fitness during sustained antigen exposure and thereby improve their infiltration, persistence, and antitumor activity in solid tumors.

## Introduction

Chimeric antigen receptor (CAR) T cell therapies have transformed hematologic cancer treatment but show limited efficacy against solid tumors^1–4^. While physical barriers, antigen heterogeneity, and suppressive microenvironments are known obstacles, overcoming them yields marginal improvements^4^, suggesting a need to re-evaluate core CAR design principles. In blood cancers, CAR Ts engage targets in a “1-on-1” or “N-on-1” mode in a highly mobile environment, allowing them a brief rest after each killing to rejuvenate^5^ ^6^. Conversely, solid tumors require but limit CAR T infiltration and force the infiltrated CAR Ts into a “1-on-N” mode of engagement^6^. Surrounded by antigen-dense tumor cells and trapped by the extracellular matrix, CAR Ts undergo intense and continuous stimulation^6^. We reason that a fast-off binding kinetics can allow CAR Ts a “break” from the intense chronic engagement in tumor mass, alleviating the activation-induced cell death (AICD) or exhaustion.

Nature has already utilized this principle for immune protection. Unlike high-affinity, slow-off monoclonal antibodies (mAbs) ^7^ ^8^, natural T-cell receptors (TCRs) have evolved to much lower affinity (*K_D_*: 1-100µM)^9^ ^10^ and fast-off kinetics (*T_1/2_*: ∼2-10s)^10–13^. Even clinically successful, affinity-enhanced TCRs preserve relatively fast-off rates. For example, the MAGE-A4 TCR that won FDA approval^14^ is low affinity (*K_D_*=5.1µM). Although its *k_off_* and *T_1/2_* are not reported, the *T_1/2_*of its parental TCR is 0.9s with *K_D_* of 65µM^15^. The NY ESO^c259^ TCR-T that generated impressive clinical efficacy is also low affinity and fast-off (*K_D_*=795nM; *T_1/2_*=19s)^16^ ^17^, establishing the dissociation rate (*k_off_*) as a robust biomarker of T cell potency^18^. In contrast, most CARs use high-affinity, slow-off antigen-binding domains^6^. A few studies found that low-affinity CARs could mitigate on-target/off-tumor toxicity^19–21^ and showed improved efficacy in preclinical^21–25^ and clinical studies that were summarized in our recent meta-analysis of solid tumor CAR T trials^6^. However, many of these investigations utilized mAbs recognizing distinct epitopes, confounding the effects of affinity with epitope geometry and/or other molecular features. Moreover, those studies have largely overlooked the binding kinetics, especially the *k_off_*, which governs antigen binding dwell time, rebinding, and cumulative signaling. Consequently, how *k_off_* shapes CAR T tumor infiltration, persistence and exhaustion in antigen-rich solid tumors remains unclear.

Here, we address this gap using a pair of EGFR-targeting mAbs, Cetuximab (Cet) vs it humanized variant H1, with matched fast association rates (*k_on_*) but markedly divergent *k_off_* ^26^, isolating antigen-binding dwell time from epitope specificity. We demonstrate that fast-on/fast-off (fast-on/off) binding reduces CAR antigen occupancy and engagement stability. Importantly, under conditions modeling limited CAR T treating large established tumors, fast-off H1 CAR Ts exhibit superior tumor infiltration, expansion, persistence, and therapeutic efficacy. Mechanistically, fast-off binding kinetics limit CAR downregulation and exhaustion, preserving CAR T metabolic and functional fitness. Our findings establish *k_off_* and the resulting antigen-binding dwell time as a critical determinant of CAR T tumor infiltration and function. Our study provides strong evidence to shift CAR design toward antigen-binding kinetics, especially the dissociation rates, to develop CAR T cells that possess enhanced tumor infiltration and functional fitness for effective solid tumor immunotherapies.

## Materials and Methods

### Mice and small-animal imaging

NOD-scid IL2Rγ^null (NSG) mice were purchased from The Jackson Laboratory. Eight- to 10-week-old mice were used. Tumor growth was monitored by bioluminescence imaging (BLI) or fluorescence imaging (FLI) using an Ami X spectral imaging system (Spectral Instruments). All animal procedures were approved by the Institutional Animal Care and Use Committee of Augusta University.

### Cell culture

A431, FaDu, A549, Jurkat, and HEK293T cells were obtained from ATCC and maintained in recommended media. Cell lines were used for no more than 12 passages. A431-Luci-iRFP720 cells were generated by lentiviral transduction with pHIV-iRFP720-E2A-Luc (Addgene). A431-GFP, FaDu-GFP, and A549-GFP cells were generated by transduction with pCDH-EF1-GFP, constructed by inserting EGFP into pCDH-EF1 (Addgene).

### CAR construction, Human CAR-T and CAR-Jurkat cells

EGFR-specific second-generation CARs were constructed using scFvs derived from paired monoclonal antibodies: Cetuximab and its humanized derivative H1 (*60*), or H528 and 2HH11 (*27*). CARs consisted of a CD8 leader, VL, a 4×GGGGS linker, VH, CD8 hinge, CD28 transmembrane domain, and 4-1BB/CD3ζ intracellular signaling domains. mCherry or NanoLuc (NanoLuc-P2A) was fused to the C-terminus of CD3ζ for CAR expression tracking. CAR cassettes were cloned downstream of the human EF1α promoter in pCDH lentiviral vectors (Addgene). Lentivirus was produced as previously described ^27^ ^28^.

Human CAR-Ts were prepared as we previously described ^24^. Briefly, T cells were isolated from healthy donor PBMCs using the EasySep Human T Cell Isolation Kit (StemCell Technologies) and activated with CD3/CD28 Dynabeads (Fisher Scientific), followed by lentiviral transduction. CAR expression was detected using recombinant human EGFR-Fc chimera (rhEGFR-AF647; R&D Systems). Jurkat cells were transduced using the same procedure without CD3/CD28 activation.

### Flow cytometry

Cells were stained with indicated antibodies and analyzed using NovoCyte Quanteon or Penteon flow cytometers (Agilent). CAR expression was assessed using mCherry or rhEGFR-AF647 together with antibodies against human CD45, CD3, CD4, and CD8. Annexin V-APC was used to assess apoptosis, and anti-PD-1 staining was used to evaluate T-cell exhaustion. For *in vivo* studies, rat anti-mouse CD45-APC was used as an internal reference.

### SPR binding kinetics

Binding kinetics of Cet and H1 scFvs were determined by surface plasmon resonance (SPR) using a Biacore 8K system (Cytiva). Recombinant Cet, H1, H528, and 2HH11 scFv-Fc proteins were produced in CHO-K1 cells and purified by protein A chromatography. Anti-human Fc antibody was immobilized on a CM5 chip, followed by capture of scFv-Fc proteins. Recombinant human EGFR extracellular domain was injected at eight concentrations (0–50 nM), and association and dissociation kinetics were analyzed using a 1:1 binding model.

### Soluble EGFR binding by CAR-T cells

mCherry-positive Cet- or H1-CAR-T cells (5×10^4^/well; triplicate) were incubated with rhEGFR-AF647 (1.6 ng/mL–25 µg/mL) for 30 min at room temperature in PBS containing PE/Cy7-conjugated anti-human CD45. Cells were washed and analyzed by flow cytometry. EGFR-binding MFI was normalized to mCherry MFI.

### CAR-Jurkat binding to EGFR-positive tumor cells

CAR-expressing Jurkat cells were evaluated for tumor-cell binding by measuring CAR-T/tumor-cell doublets as previously described ^29^. mCherry-positive CAR-Jurkat cells (5×10^4^) were incubated with A431-GFP cells at indicated ratios for 5 min at 4°C followed by 30 min at either 4°C or 37°C. After washing, cells were analyzed by flow cytometry. Binding capacity was defined as the percentage of mCherry^+GFP^+ double-positive cells among total mCherry^+ cells.

### CAR-T proliferation and cytotoxicity

For tumor-induced proliferation, A431-GFP cells (1×0^6^/well) were cultured overnight in 6-well plates. CAR-T cells were labeled with CellTrace Violet (Invitrogen) according to the manufacturer’s instructions and cocultured with tumor cells as previously described ^30^. The number after proliferation is compared to original cells to calculate the fold of expansion.

For real-time cytotoxicity assays, A431-GFP cells (1×10^4^/well; triplicate) were seeded in 96-well plates overnight and cocultured with CAR-T cells at effector-to-target ratios of 0.5:1, 1:1, or 2:1. Images were acquired every 4–6 h for 3–5 days using an IncuCyte live-cell imaging system (Sartorius). Cytotoxicity was quantified by measuring GFP-positive tumor-cell area.

### Apoptosis Staining

CAR-T apoptosis was assessed by Annexin V staining according to the manufacturer’s protocol. Live cells were defined as human CD45^+^mCherry^+^Annexin V^−^ and apoptotic cells as human CD45^+^mCherry^+^Annexin V^+^.

### Adoptive Transfer and Monitoring of CAR T Cells

Ten to 12 days post-lentiviral transduction, EGFR CAR T cells were adoptively transferred into tumor-bearing NSG mice via tail vein injection. Circulating CAR T cells were monitored in peripheral blood by staining with anti-human CD45, CD4 or CD8 antibodies, together with mCherry fluorescence (for BBz-mCh CAR T cells) or Recombinant Human EGFR Fc Chimera-AF647 (for BBz-Nluc CART cells). Mouse CD45-positive leukocytes were used as an internal reference for normalization.

For NanoLuc imaging, fluorofurimazine (FFz), a high-output NanoLuc substrate (*64*), was prepared in poloxamer-407 (Pluronic F127; Sigma-Aldrich) according to the published protocol. Tumor burden was monitored by FLI of A431-Luci-iRFP720 cells, whereas NanoLuc-expressing CAR-T cells were quantified by BLI using the Ami X system.

### Tumor-infiltrating lymphocytes and immunohistochemistry

Tumors were collected, washed with PBS, mechanically dissociated, and digested for 30 min at 37°C in serum-free RPMI-1640 containing 0.1% collagenase V, 0.1% hyaluronidase, and 100 U/mL DNase I. Following red blood cell lysis, single-cell suspensions were analyzed by flow cytometry and functional assays.

For immunohistochemistry, tumors were fixed in 10% buffered formalin for 24 h, transferred to 70% ethanol, paraffin embedded, sectioned, and stained with rabbit anti-human CD8 antibody followed by HRP-conjugated anti-rabbit IgG and TMB visualization.

### Single-cell RNA sequencing

CAR-T cells were collected after 2 days of *in vitro* A431 stimulation or from mouse spleens on days 14 and 20 after adoptive transfer. Viable cells were enriched using MojoSort Human or Mouse Dead Cell Removal Kits (BioLegend). Single-cell suspensions were processed using the 10X Genomics Chromium X platform with Single Cell 3′ Gene Expression v4 reagents. Samples were multiplexed and sequenced as separate libraries. Raw data were processed using Cell Ranger v10 with combined human (GRCh38) and mouse (GRCm39) reference genomes supplemented with CAR-vector-specific WPRE-mCherry sequences. Downstream analyses were performed as previously described ^31^.

### Statistical analysis

Statistical analyses were performed using GraphPad Prism 10. Two-tailed Student’s *t*-test or two-way ANOVA was used as appropriate. Additional statistical methods are indicated in the corresponding figure legends. Data are presented as described in figure legends.

## Results

### 1. Identification of paired EGFR mAbs with similar *k_on_* but divergent *k_off_* (Fast-off vs Slow-off)

To identify paired mAbs that bind the same epitope, but differ primarily in dissociation kinetics (*k_off_*), we selected two pairs of EGFR mAbs based on their reported *k_on_* and *k_off_*, i.e., Cet vs its humanized derivative H1^26^ and h528 vs its affinity-enhanced variant 2HH11^32^, which fit this slow-off vs fast-off criteria (table S1). Because the kinetic parameters vary widely among different reports ^26^ ^33–36^ (table S2) and our objective is to identify fast-off binders for CAR construction, we re-evaluated their binding kinetics in scFv-Fc format using surface plasmon resonance (SPR). The data showed Cet and H1 exhibited similarly rapid association kinetics (*k*_on_=7.81 and 9.41×10□M□¹s□¹, respectively), indicating comparable fast-on rates. In contrast, their *k_off_* diverged substantially. The sensorgrams demonstrated sustained binding of Cet during the dissociation phase (*k*_off_=1.52×10^-3^ s^-1^; T_1/2_ =456s), whereas H1 rapidly disengages following analyte removal (*k*_off_=2.77×10^-2^ s^-1^; T_1/2_ =25s) (Fig. 1A-B). This pronounced 18-fold difference in *k*_off_ establishes a clear dwell time divergence between Cet and H1. Additionally, H1 exhibited reduced relative binding response compared to Cet (58.43% vs 71.38%), suggesting reduced analyte engagement under identical capture conditions. In contrast, the pair of h528 and 2HH11 mAbs displayed nearly identical *k_on_* and only minor differences in *k_off_*(fig. S1A-B), resulting in just 2-fold difference in dwell time, sharply different from a prior report^32^.

**Fig. 1.**
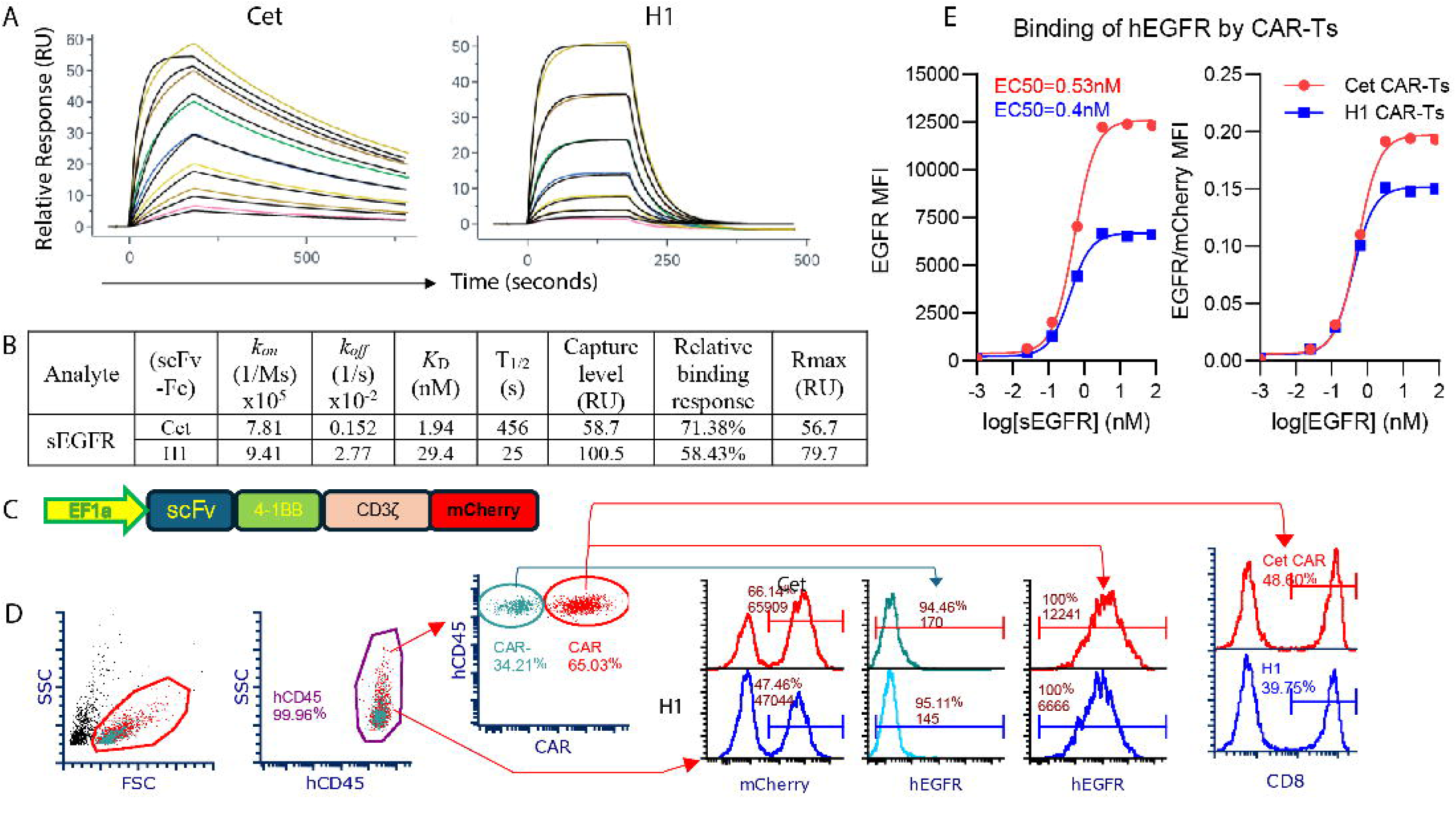
Fast-off kinetics limit steady-state EGFR engagement by scFv antibodies and CAR T cells. **(A)** SPR sensorgrams depicting the binding kinetics of soluble hEGFR (sEGFR) to captured scFv-Fc (Cet and H1). Black lines represent global kinetic fits using a 1:1 Langmuir binding model. **(B)** Summary table of biophysical parameters derived from SPR analysis, including association rate constant (*k*_on_), dissociation rate constant (*k*_off_), equilibrium dissociation constant (*K*_D_), dissociation half-life (*T*_1/2_), capture level, relative binding response, and maximum binding capacity (*R*_max_). **(C)** Schematic diagram of the second-generation 4-1BB-based chimeric antigen receptor (CAR) construct containing an mCherry reporter sequence downstream of the CD3ζ signaling domain. **(D)** Flow cytometry gating strategy evaluating CAR expression (mCherry^+^), the % of CD8 in the CAR T cells, and the soluble hEGFR binding capacity on gated CAR^-^ and CAR^+^ T cell populations. Representative histograms display mCherry mean fluorescence intensity (MFI) and hEGFR binding profiles for Cet- and H1-CAR T cells. **(E)** Dose-response binding curves of soluble hEGFR to Cet- and H1-CAR T cells, displaying the raw EGFR MFI (left) and the normalized EGFR/mCherry MFI ratio (right) across a range of log[sEGFR] concentrations. Calculated EC_50_ values are indicated. Data are representative of three independent experiments with similar results.

While SPR measures binding kinetics in a controlled biochemical setting, CAR molecules engage antigen within a two-dimensional cell membrane interface. To study how the *k*_off_ divergence between Cet and H1 in soluble scFv-Fc format would be preserved when expressed as CARs on T cells, we stained CAR Ts with soluble hEGFR and analyzed them by flow cytometry. To control potential differences in CAR expression, Cet- and H1-based CAR constructs were linked to the mCherry reporter, thereby enabling normalization of EGFR binding to CAR levels (Fig. 1C). The data showed that H1 CAR T exhibited a substantially lower maximal EGFR binding plateau compared to Cet CAR T (Fig. 1D-E). This reduced antigen occupancy is consistent with the fast-off rate of H1 observed by SPR, indicating that its rapid dissociation limits sustained antigen engagement in the membrane-bound CAR context. Together, the findings demonstrate that intrinsic molecular dwell time is preserved as a functional determinant of antigen occupancy upon incorporation into CAR. Consistent with SPR data, h528 and 2HH11 CAR Ts displayed nearly superimposable EGFR binding curves, both in absolute MFI and after normalization to CAR expression (fig. S1C-D).

Collectively, we have identified the Cet/H1 pair as a robust kinetic contrast suitable for interrogating how dissociation kinetics will affect CAR T function. Accordingly, subsequent studies were conducted using the Cet-/H1-based CAR constructs to interrogate how *k*_off_ affects CAR T’s tumor infiltration, function, persistence, and antitumor effects.

### 2. Fast-off H1 CAR T/tumor cell engagement is unstable and causes less CAR stripping

To evaluate CAR T/tumor engagement, we analyzed CAR T/tumor cells doublet formation and CAR loss on T cells after a brief 30min co-culture. The data showed that fast-off H1 CAR Ts formed fewer doublets with A431 tumor cells. In addition, H1 CAR T cells after binding target tumor cells caused less CAR loss at lower E:T ratios (fig. S2). Since the CAR T/A431 cell physical interaction is complicated by the CAR T’s cytotoxicity on target cells. We thus studied the physical interaction between CAR-Jurkat and tumor cells (Fig. 2A). At both 4°C and 37°C, slow-off Cet CAR-Jurkat demonstrated more doublet formation (Fig. 2B-F). Notably, shifting the co-culture to 37°C revealed a pronounced difference in engagement stability: H1 CAR-Jurkat/A431 cell doublets dissociated at a significantly higher rate than their Cet counterparts (Fig. 2G). Furthermore, this faster dissociation was accompanied by better maintenance of CAR expression. Following separation from tumor cells, Cet CAR-Jurkats exhibited a nearly 50% reduction in CAR mean fluorescence intensity (MFI) at low E:T ratios, whereas H1 CAR-Jurkats retained most of their receptors, showing only ∼20% decrease (Fig. 2H). The CAR downregulation was driven by bidirectional trogocytosis at the immunological synapse. Tumor cells co-cultured with Cet CAR-Jurkat exhibited a profound increase in mCherry (Fig. 2I), indicating active receptor extraction by target cells. Reciprocally, Cet CAR-Jurkat demonstrated more acquisition of tumor-derived GFP (Fig. 2J). Taken together, these data indicate that the fast-off H1 CAR Ts form a more transient, unstable interaction that helps preserve surface CAR density. Conversely, the slow-off Cet CAR T with prolonged engagement exacerbates bidirectional trogocytosis, prompting severe CAR stripping and depletion upon cell dissociation.

**Fig. 2.**
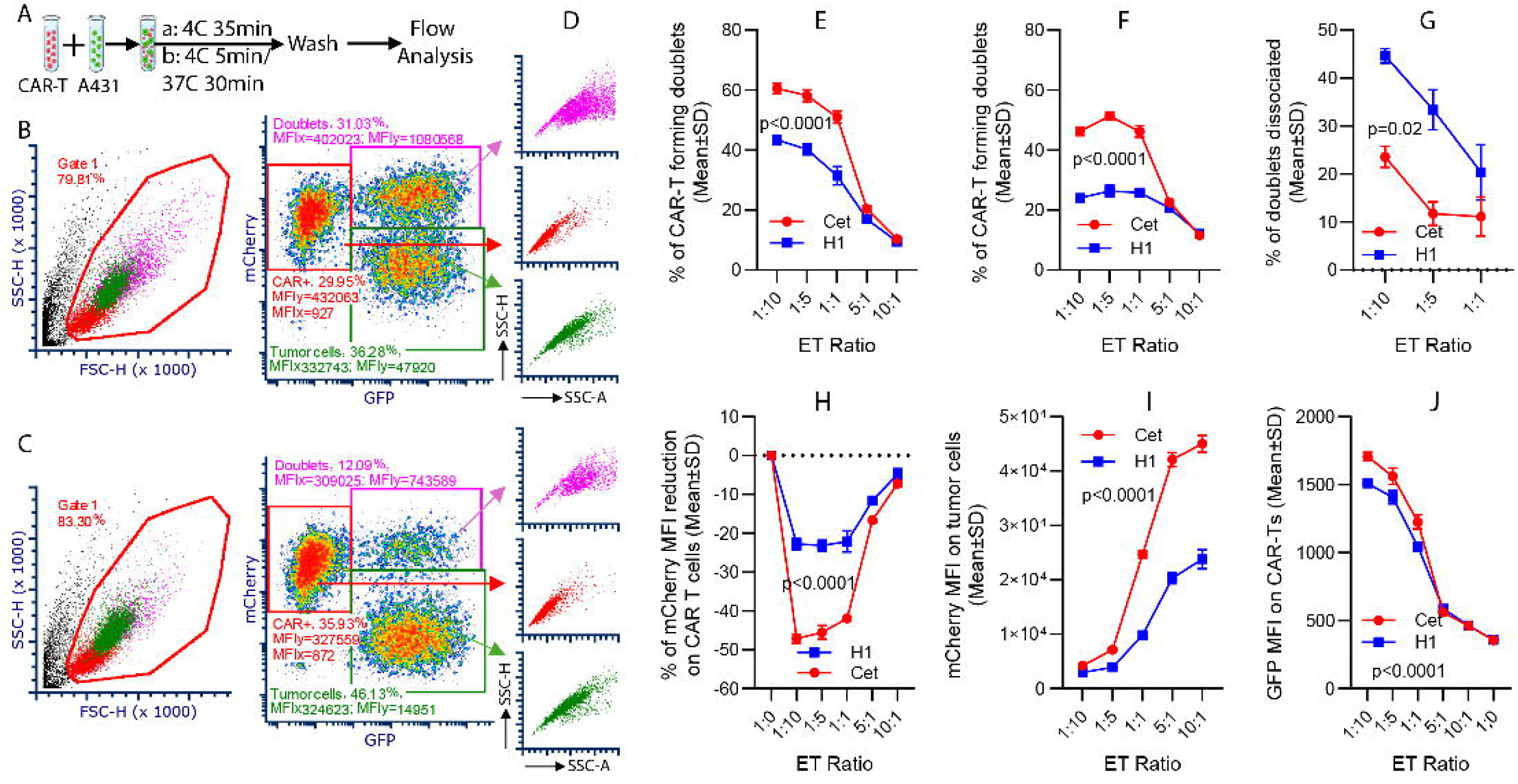
Fast-off H1 CAR-Jurkat cells exhibit transient engagement stability and resist bidirectional trogocytosis-mediated CAR stripping. (**A**) Experimental workflow schema. Cet or H1 CAR-Jurkat cells (red) were co-cultured with A431 target tumor cells (green) at the indicated effector-to-target (E:T) ratios at 4°C for 5 min, followed by a 30-min incubation at either 4°C or 37°C prior to flow cytometric analysis. (**B and C**) Representative flow cytometry plots and gating strategies identifying cell doublets (magenta gate) formed by Cet **(B)** or H1 **(C)** CAR-Jurkat cells and tumor cells. **(D)** Backgating analysis showing the side scatter H (SSC-H) vs side scatter area (SSC-A) profiles of the isolated doublet populations. **(E and F)** Quantification of CAR-Jurkat/tumor cell doublet percentages following incubation at 4°C **(E)** or after shifting to 37°C **(F)** across various E:T ratios. **(G)** Percentage of doublets dissociated during the 37°C incubation phase, calculated by comparing doublet frequencies at 4°C versus 4°C-to-37°C conditions. **(H)** Relative percentage reduction of CAR level (mCherry MFI) on dissociated CAR-Jurkat cells (cells in the red square). **(I and J)** Evaluation of bidirectional trogocytosis at specified E:T ratios. **(I)** Transfer of CAR molecules (mCherry MFI) onto the surface of gated target tumor cells (cells in green square). **(J)** Acquisition of tumor-derived GFP (GFP MFI) by gated CAR-Jurkat cells (cells in the red square). Data are presented as mean ± SD from triplicates. Statistical significance was determined by two-way ANOVA (p-values indicated on plots). Data are representative of two independent experiments.

### 3. Fast-off H1 CAR Ts accumulate more than slow-off Cet CAR Ts after *in vitro* engagement with target cells despite comparable proliferation and cytotoxicity

To compare how CARs with distinct *k_off_* respond upon target antigen encounter, we co-cultured H1 and Cet CAR Ts with EGFR^high^ A431 or EGFR^moderate^ FaDu target cells (Fig. 3A). Both CAR Ts mediated comparable cytotoxicity against A431 and FaDu cells at a high E/T ratio, but H1 CAR T showed stronger killing activity at a lower E/T ratio (Fig. 3A-B). H1 and Cet CAR Ts underwent comparable rounds of proliferation following tumor engagement, but the magnitude of each division peak was higher for H1 CAR Ts (Fig. 3C-D), indicating more accumulation of expanding H1 CAR T cells. Indeed, the absolute number and thus the expansion folds of H1 CAR Ts after co-culture with A431 tumor cells was significantly higher than that of Cet CAR Ts (Fig. 3E). The enhanced accumulation of H1 CAR Ts following co-culture with EGFR^moderate^ FaDu cells was less obvious (fig. S3). Together, the results indicate that under high antigen stimulation, the fast-off H1 CAR Ts could accumulate more, and that the enhanced accumulation of H1 CAR Ts is not due to increased proliferative capacity but likely reflects lower AICD and improved survival following antigen engagement.

**Fig. 3.**
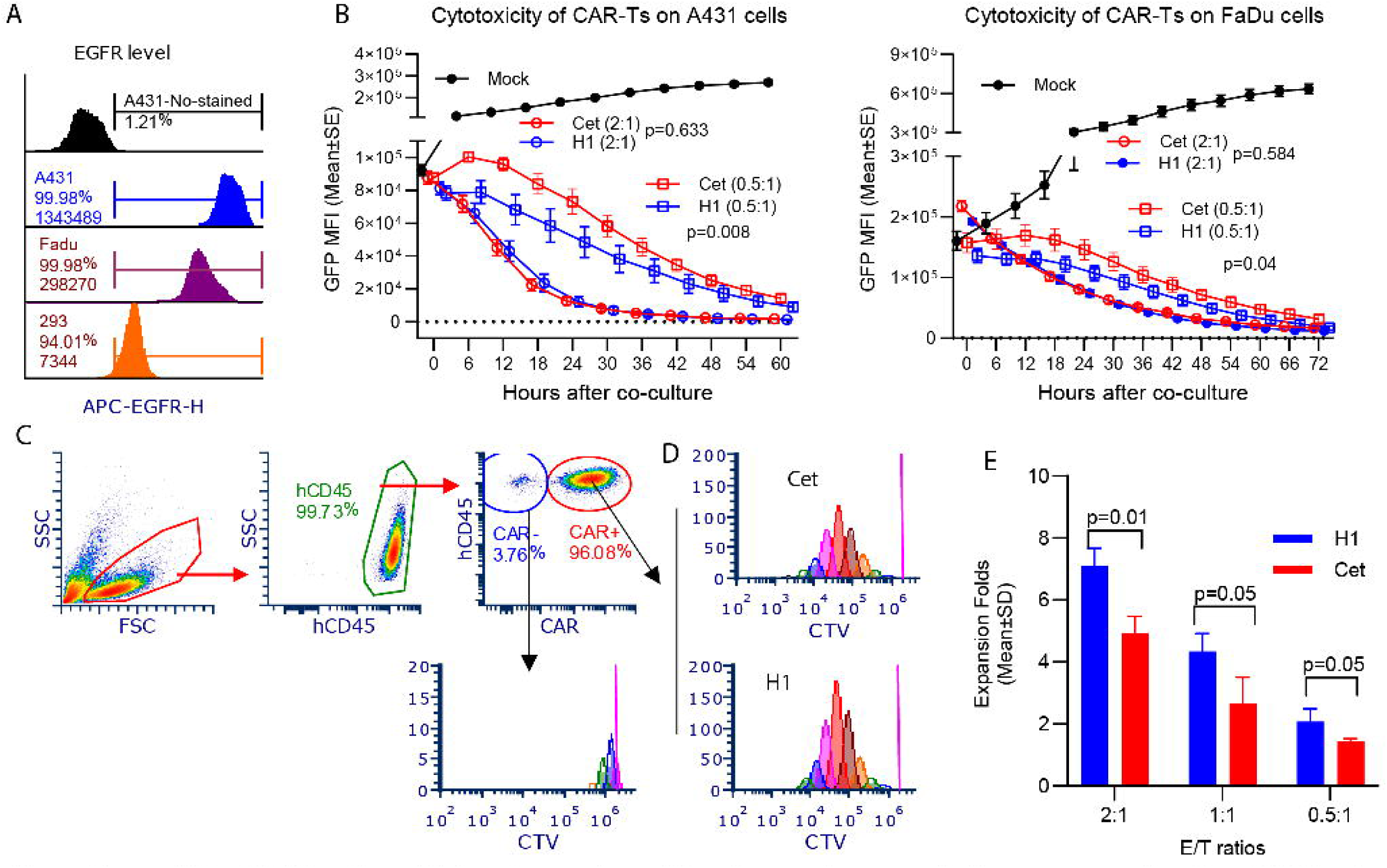
Fast-off H1 CAR T cells exhibit enhanced proliferation and accumulation compared to Cet CAR T cells upon engagement with EGFR^+^ tumor cells. (**A**) Flow cytometry histograms showing surface EGFR levels on A431, FaDu, and HEK293 cells, stained with anti-EGFR antibody. Non-stained A431 cells served as a negative control. (**B**) Real-time cytotoxicity kinetics of Cet and H1 CAR T cells co-cultured with GFP^+^ A431 (left) or FaDu (right) tumor cells at indicated E:T ratios, tracked by continuous monitoring of target cell GFP using an IncuCyte live-cell imager. (**C**) Gating strategy defining live, hCD45^+^ T cells, further segregated into CAR^-^(blue gate) and CAR^+^ (red gate) populations to evaluate longitudinal CAR T cell expansion and proliferation. (**D**) Flow cytometric tracking of CAR T cell proliferation via CellTrace Violet (CTV) dye dilution after tumor engagement. Successive peaks represent cell divisions, with the peak amplitude corresponding to cell numbers. The non-dividing, un-stimulated CAR^-^T cell population (magenta vertical line/peak) serves as the baseline control. **(E)** Fold-expansion of CAR T cells calculated relative to the initial number of input T cells following co-culture with target cells across the indicated E:T ratios. Data are representative of three independent experiments and are presented as mean ± SD or mean ± SE as indicated on axes. Statistical significance was determined by two-way ANOVA **(B)** or unpaired Student’s t-test **(E)** (p-values are indicated within the panels).

### 4. Cet and H1 CAR Ts exhibit comparable antitumor efficacy at high doses

To compare the *in vivo* antitumor activity of Cet and H1 CAR Ts, we evaluated their therapeutic efficacy using the A431 xenograft tumor model. In the high-dose cohort, mice were inoculated with 1×10 A431-Luc-iRFP720 tumor cells (A431-Luc/720), followed by the adoptive transfer of 5×10 CAR Ts on day 8 after tumor implantation. Under these conditions, both Cet and H1 CAR Ts completely eradicated established tumors, and all treated mice remained tumor-free for the duration of the study (Fig. 4A-B). Similar numbers of Cet and H1 CAR Ts were detected in mouse blood (Fig. 4C, fig. S4). The tumor size data at the end of experiments are summarized in Fig. 4D. The mice treated with Cet CAR Ts had a slight (but statistically insignificant) survival disadvantage (Fig. 4E). These findings indicate that, at a high cell dose, Cet and H1 CAR Ts exhibit comparable antitumor efficacy.

**Fig. 4.**
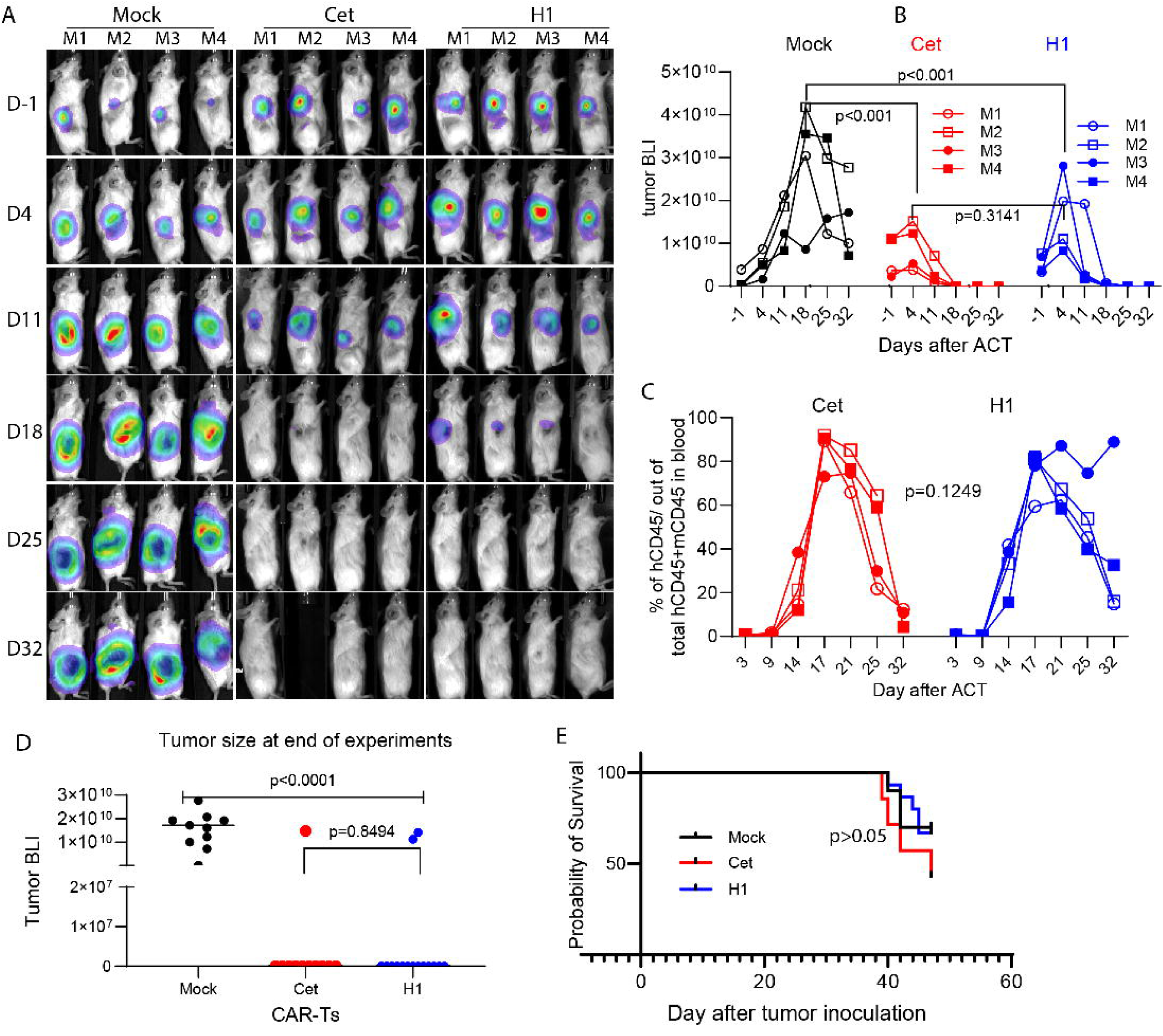
High-dose fast-off H1 and slow-off Cet CAR T cells exhibit comparable antitumor efficacy and *in vivo* expansion in a small-tumor model. NSG mice were inoculated with 1×10^6^ A431-Luc/720 tumor cells and treated on day 8 with 5×10^6^ CAR T cells. **(A)** Longitudinal tumor BLI of individual mice (M1–M4) across treatment groups at indicated time points relative to adoptive cell transfer (ACT; Day -1 to Day 32). **(B)** *In vivo* tumor progression kinetics quantified by BLI total flux for individual mice over time. The decline in BLI signal observed in the mock group at later time points is attributed to limited luciferase substrate penetration within large tumor masses. **(C)** Longitudinal tracking of primary human T cell in peripheral blood, expressed as the percentage of hCD45^+^ cells within total leukocytes (hCD45^+^ + mCD45^+^ [mouse CD45^+^]). Individual mouse symbols correspond to definitions in **(B)**. Gating strategy is detailed in Figure S4. **(D)** Statistical summary of terminal tumor burden (BLI) at the experimental endpoint, pooled from three independent experiments (n=12-15 mice per group). **(E)** Kaplan-Meier survival curves of tumor-bearing mice across the indicated treatment cohorts. Data are representative of three independent experiments. Statistical significance was determined by two-way ANOVA **(B and C)** or unpaired Student’s t-test **(D)** (p-values are indicated within the panels).

### 5. Treatment of large tumors with low dose T cells reveals superior expansion, function, and therapeutic efficacy of fat-off H1 CAR T

To simulate the clinical scenario that large tumors are often treated with limited numbers of CAR Ts, we increased the tumor burden by inoculating 2×10 A431-Luc/720 cells and reduced the CAR T dose (3×10 cells). Since initial tumor burden is relevant to therapeutic outcome, we divided the tumor bearing mice to make sure that the Cet and H1 treated groups had similar iRFP720 MFI when starting treatment (fig. S5A-a). Under this condition, the therapeutic differences between Cet and H1 CAR Ts were evident. The H1 CAR T was able to mediate significantly stronger tumor control (p=0.018) than Cet CAR T, and even regression in a subset of mice (Fig. 5A-B, fig. S5A-a). The strong efficacy of H1 CAR Ts correlated with superior *in vivo* expansion at day 13 time point after therapy and then re-expanded after day 19 (Fig. 5C, fig. S5A-b/c). In contrast, Cet CAR T failed to expand noticeably in 6 of 8 mice (Fig. S5A-c). This is drastically different from the comparable significant expansion of both Cet and H1 CAR Ts in treating small tumors with high-dose cells (Fig. 4C). Even in the mouse with significant initial detection of Cet CAR T cells (Cet-M1), the expansion was short-lived, and tumor control was transient (Fig. 5D-E). In contrast, the two mice (M2 and 4) in H1 group nearly eradicated the tumors, in which the H1 CAR T cells underwent re-expansion from day 19 (Fig. S5A-c/d). The initial re-expansion from day 19 was likely driven by antigen as the % of CAR+ and % of CD8 T cells remained high (Fig. S5A-e). However, the re-expansion after Day 27 might be complicated by GvHD, reflected by the percentage reduction of CAR and CD8 T cells (Fig. S5A-e). Nevertheless, our data show that, in large tumors, fast-off H1 CAR Ts demonstrated stronger antitumor effects with greater initial antigen-driven expansion, with higher potential of re-expansion after further encounter with tumor antigen.

**Fig. 5.**
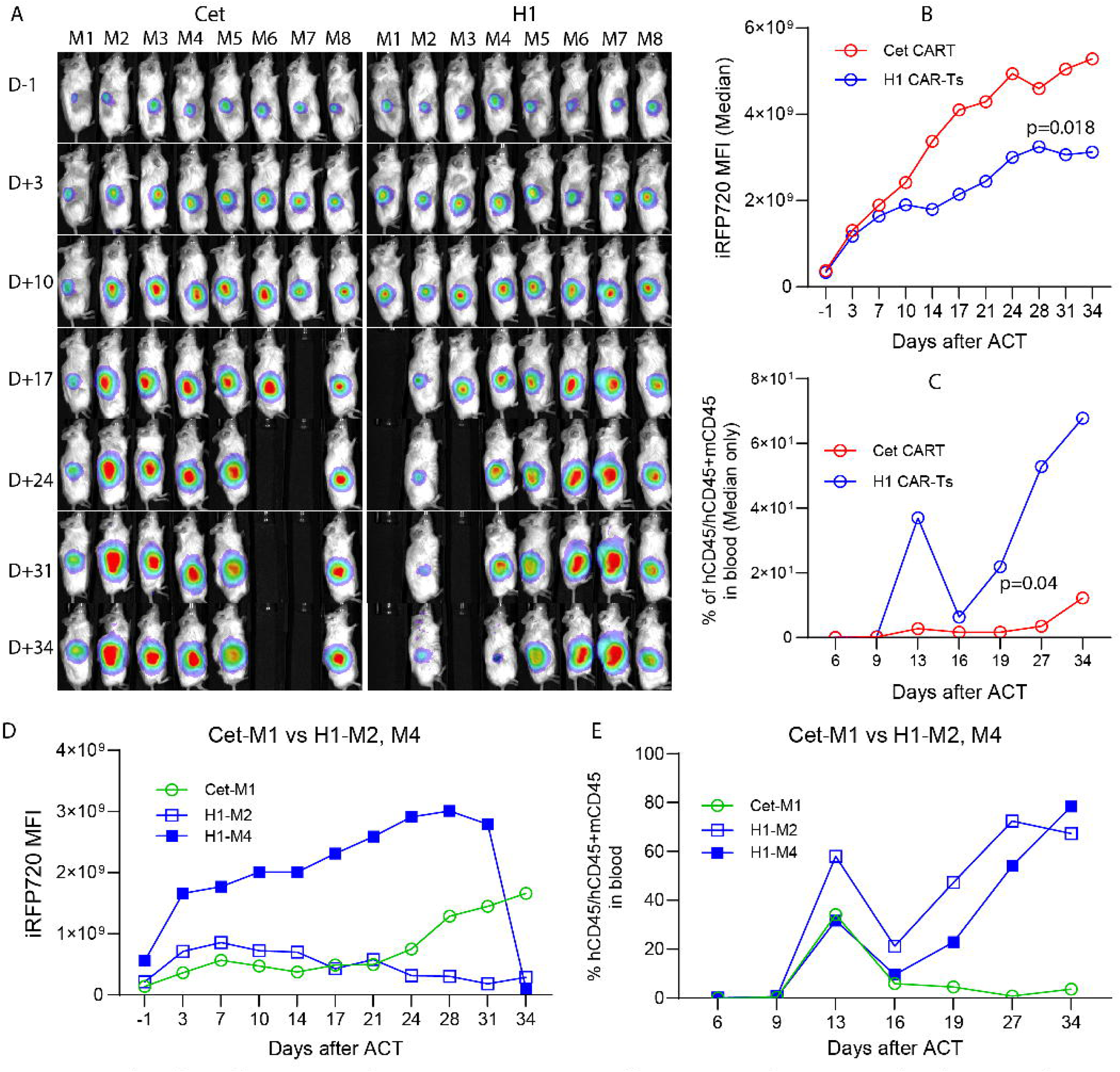
Low-dose fast-off H1 CAR T cells elicit superior antitumor efficacy against large tumor burdens, correlating with enhanced *in vivo* expansion. NSG mice were inoculated with 2×10^6^ A431-Luc/720 tumor cells and treated on day 8 with 3×10^6^ CAR T cells. **(A)** Longitudinal iRFP720 imaging tracking large solid tumor burdens in individual mice (M1–M8) across indicated time points relative to adoptive cell transfer (ACT; Day -1 to Day 34). Infrared 720 imaging was utilized to circumvent the tissue penetration and substrate limitations of bioluminescence imaging within large tumor masses. (**B and C)** Longitudinal median tumor iRFP720 MFI **(B)** and the median percentage of human T cell (% hCD45□out of total human + mouse CD45□cells) in peripheral blood **(C)** over time (*n* = 6–8 mice per group; gating strategy outlined in Supplemental Figure S4). **(D and E)** Longitudinal single-mouse kinetic trajectories directly comparing tumor progression (iRFP720 MFI) **(D)** and peripheral blood hCD45 **(E)** for selected individual animals (Cet-M1 and M5 versus H1-M2 and M4). Note: Complete tumor regression followed by eschar detachment and total dermal wound healing at the initial injection site was observed in the H1-M4 mouse by day 34 (D34). Data are representative of three independent experiments with similar results. Statistical significance for median kinetic curves was determined by the Wilcoxon signed-rank test (*p-*values are indicated on the plots).

To further analyze the CAR T antitumor effects and expansion *in vivo*, we also constructed the Cet and H1 CAR Ts with co-expression of nanoluciferase, which allowed us to monitor CAR T expansion in the whole body. Under the same setting of treating large tumors with low-dose T cells, H1 CAR T again generated stronger antitumor effects than Cet CAR T (Fig. S5B). The blood CAR T and whole body nanoluciferase activity demonstrated that H1 CAR Ts generated more expansion than Cet CAR T, correlated to better antitumor effects.

### 6. H1 CAR T demonstrated superior tumor infiltration, function, and immune memory

We evaluated the spatiotemporal CAR T infiltration by immunohistochemical staining (IHC) of the tumors harvested on days 14, 18, and 20 after treatment. On day 14, Cet and H1 CAR Ts were detected at the tumor periphery, and the H1 CAR T clusters were dense and larger (fig. S6A-a). However, by day 18 and 20, a clear divergence in spatial distribution and density emerged. While the Cet CAR T became less frequent and remained sequestered at the tumor edge (Fig. 6A), H1 CAR T demonstrated markedly deeper penetration into tumor parenchyma (Fig. 6B). Strikingly, large intratumoral T-cell clusters were observed exclusively in the H1 group, indicating robust local expansion that slow-off Cet CAR T failed to achieve. This was further confirmed by flow analysis showing significantly more H1 CAR T accumulation in tumor compared to Cet CAR T (Fig. 6C-D) even though the Cet CAR T counts in the spleen is higher. The observation of deeper tumor infiltration and larger H1 CAR T clusters is highly reproducible, being observed in all three tumors examined (two mice on D18 and one mouse on D20). This strongly suggests that the frequent disengagement during pulsatile on/off binding frees fast-off CAR Ts to avoid entrapment at tumor edge and to migrate deep into tumor.

**Fig. 6.**
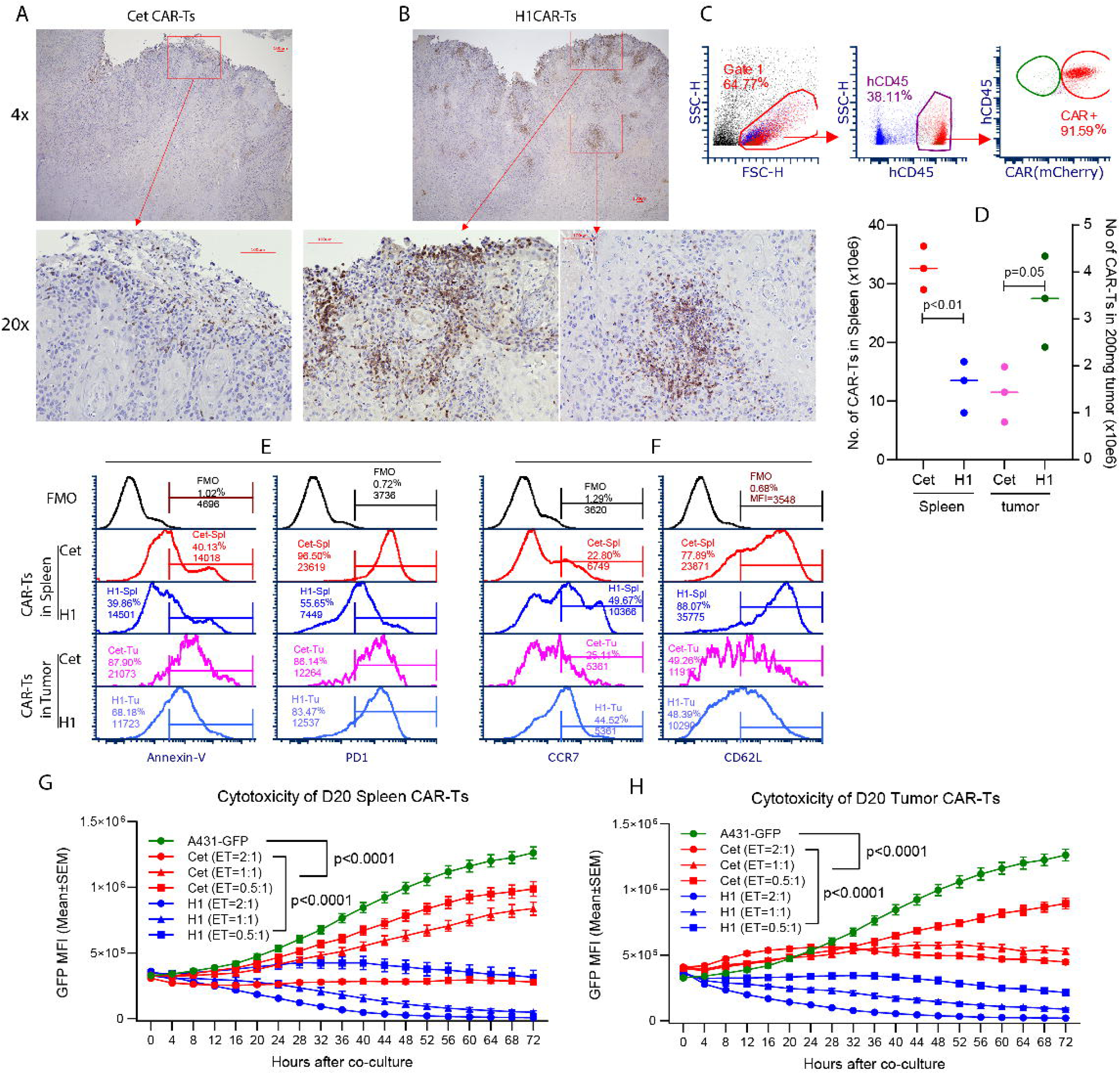
Fast-off H1 CAR T cells exhibit deeper tumor infiltration, enhanced accumulation, reduced exhaustion, and superior therapeutic efficacy compared to slow-off Cet CAR T cells. **(A and B)** IHC staining evaluating CAR T cell tumor infiltration 20 days after treatment with slow-off Cet (A) or fast-off H1 (B) CAR T cells. Representative low-magnification (4x, top panels) and high-magnification (20x, bottom panels) fields are displayed; red inset boxes in the upper panels delineate the corresponding regions of interest magnified below. **(C-F)** Spleen and tumor single cell suspension were prepared from the treated mice. The CAR T cells were analyzed by flow cytometry (C). Absolute CAR T cell numbers quantified within the spleen (left axis) and per 200 mg of isolated tumor tissue (right axis) (D). Statistical significance was determined by an unpaired Student’s t-test. The cytometric histogram profiles of CAR T cells from the spleen and tumor lesions assessing apoptosis (Annexin-V), exhaustion (PD-1 (E), and memory phenotypes (CCR7 and CD62L) (F). Fluorescence minus one (FMO) controls are indicated in black at the top of each stack. (G-H) *Ex vivo* real-time cytotoxicity kinetics evaluating the tumor-killing capacity of spleen-derived (G) and tumor-infiltrating (H) Cet and H1 CAR T cells against GFP^+^ A431 target cells. Data are representative of two independent experiments and are presented as mean ± SEM. Statistical significance was determined by two-way ANOVA (p-values are indicated within the panels).

To compare the function of tumor infiltrating and systemic Cet and H1 CAR Ts, we recovered them from tumor and spleen 14 Days and 20 days after treatment. On Day 14, both Cet and H1 maintained similar PD1 expression levels and tumor-killing activity (fig. S6A-b/c). However, on Day 20 after treatment, H1 CAR Ts were less exhausted and apoptotic (Fig. 6E) and maintained higher expression of central memory markers (CCR7 and CD62L) (Fig. 6F). This was especially true for the cells recovered from the spleen. This memory-like profile likely relates to the significant late-stage rebound of H1 CAR T observed in the treated mice (fig. S5A-c/d). Furthermore, H1 CAR Ts in the treated mouse spleen and tumor maintain much higher tumor killing activity than that of Cet CAR Ts (Fig. 6G-H). Similar phenomena were also observed on day 18 after treatment (fig. S6B).

Our data also revealed that the tumor-infiltrating Cet CAR T cells exhibited substantial CAR downregulation, ∼70% reduction compared to their splenic counterparts. In contrast, H1 CAR Ts exhibited only a modest ∼20% reduction in the tumor (fig. S6C). This *in vivo* CAR downregulation mirrors the CAR loss observed *in vitro* (fig. S2C), but is more pronounced, particularly in slow-off Cet CAR T cells. Moreover, upon overnight *ex vivo* culture, H1 CAR expression rapidly recovered to baseline or higher, whereas Cet CAR expression remained ∼40% lower than that of splenic CAR Ts. The impaired recovery of Cet CAR expression likely contributes to their diminished expansion and compromised antitumor efficacy. Together, these data provide strong evidence that fast-off kinetics play a key role in CAR T tumor infiltration, survival/expansion, and function maintenance to achieve strong therapeutic efficacy in solid tumors.

### 7. scRNAseq reveals that H1 CAR T maintains function, metabolic fitness, and memory phenotypes *in vivo*

To further characterize the Cet and H1 CAR Ts, we performed scRNA-seq analyses using CAR Ts recovered from *in vitro* stimulation and *in vivo* treatment. For the *in vitro* study, CAR Ts were co-cultured with A431 cells at an E:T ratio of 0.5:1 for 2 days. CAR Ts were enriched by removing dead cells. The *in vivo* CAR Ts were isolated from NSG mouse spleens on days 14 and 20 after treatment. The scRNAseq datasets revealed several notable findings (Fig. 7 and fig. S7): **1)** Following 2 days *in vitro* stimulation, Cet and H1 CAR Ts displayed highly similar gene expression profiles (fig. S7A); **2)** *In vivo*, the phenotypes of Cet and H1 CAR T cells remained broadly similar on day 14 (Fig. 7B-C), but H1 cells were actually more activated, with higher *GZMB*, *IFNG*, *TNF*, *LAG3*, *HAVCR2*, and *CD69* expression (fig. S7C); **3)** By day 20, striking differences emerged. H1 CAR T maintained a *TCF7*^+^/*CCR7*^+^/*IL7R*^+^ stem-like memory program (Fig. 7D and fig. S7A-B), consistent with the flow data in Fig. 6D, while Cet T cells drove toward terminal differentiation (*EOMES*^+^/*TOX2*^+^/*HAVCR2*^+^ with high effector output but no self-renewal) (Fig. 7D, fig. S7C). **4)** Metabolically, there was a dramatic reversal between D14 and D20 (Fig. 7E-F and fig. S7D). At D14, compared to Cet CAR T cells, H1 CAR T cells were more active in glycolysis, consistent with active *in vivo* expansion and effector function. However, by D20, the metabolic profiles was shifted. Cet cells exhibit a metabolic program characterized by increased aerobic glycolysis (the Warburg effect), a hallmark of highly activated effector T cells associated with terminal differentiation and reduced memory potential^37^ ^38^. In contrast, H1 cells preferentially utilize oxidative phosphorylation (OXPHOS), a metabolic state strongly associated with memory T-cell formation, longevity, self-renewal, and enhanced antitumor persistence^39–41^.

**Fig. 7.**
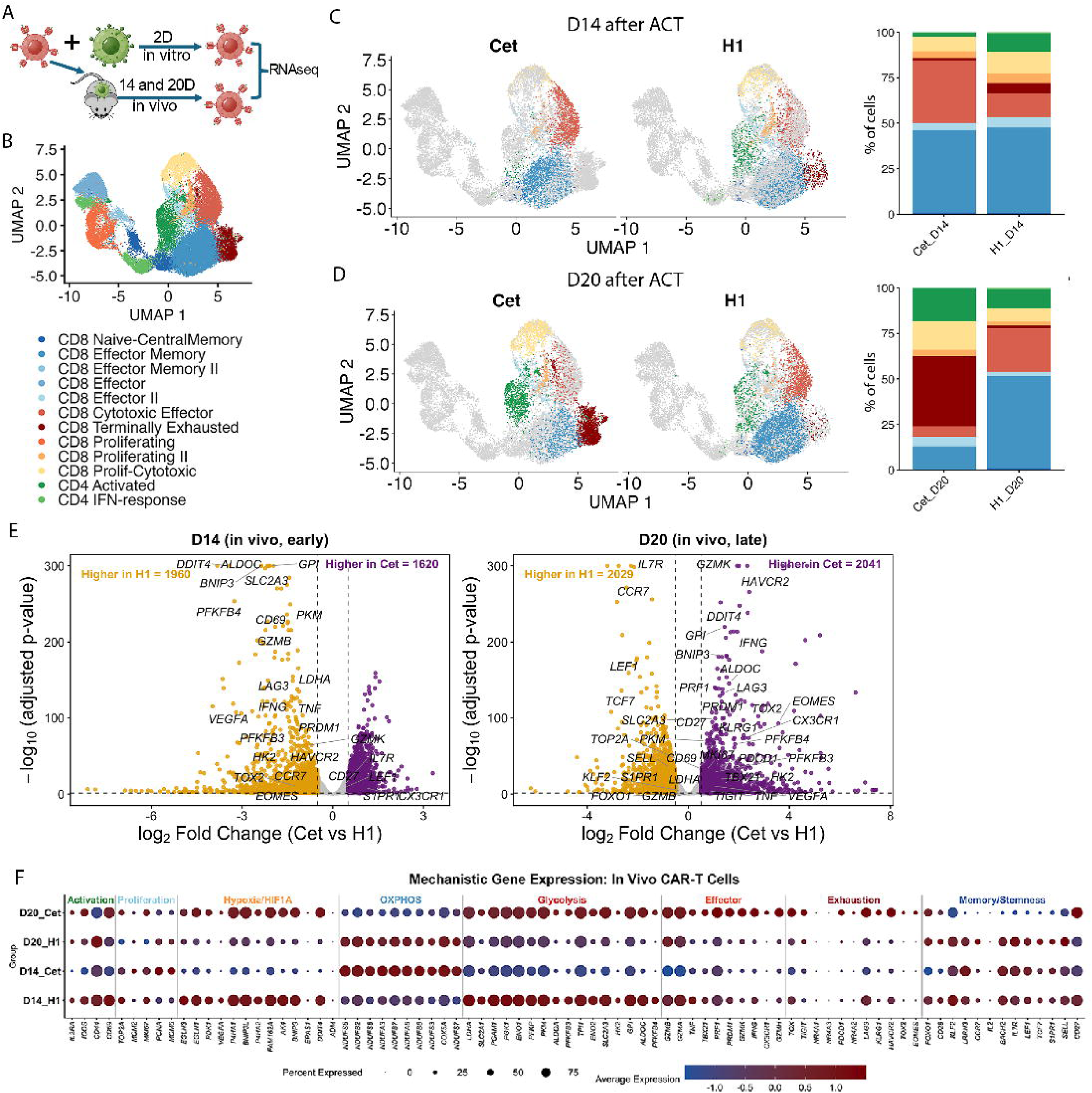
Fast-off H1 CAR T cells maintain a memory/stemness phenotype and preserve effector function *in vivo*, correlating with enhanced metabolic fitness. **(A)** Experimental design schema. CAR T cells were either co-cultured *in vitro* with A431 cells for 2 days or adoptively transferred into tumor-bearing mice and isolated from the spleen on days 14 (D14) and 20 (D20) post-injection for scRNA-seq analysis. **(B)** UMAP projection of merged single-cell data highlighting the annotated cell differentiation states and functional populations. **(C and D)** UMAP projections mapping *in vivo* Cet (slow-off) and H1 (fast-off) CAR T cells at D14 **(C)** and D20 **(D)** after ACT. Bar plots (right) summarize the relative percentage breakdown of each cell population. **(E)** Volcano plots displaying differentially expressed genes (DEGs) between fast-off H1 (gold, left) and slow-off Cet (purple, right) CAR T cells isolated at early (D14, left) and late (D20, right) *in vivo* time points. Dashed lines indicate statistical significance thresholds. **(F)** Dot plot illustrating the average expression level (color scale) and percentage of expressing cells (dot size) for mechanistic marker genes within D14 and D20 Cet and H1 CAR T cell populations, categorized by activation, proliferation, hypoxia/HIF1A, OXPHOS, glycolysis, effector, exhaustion, and memory/stemness pathways.

In summary, our findings strongly indicate that under conditions of limited CAR T cell input and high tumor burden, compared to slow-off Cet CAR Ts, fast-off H1 CAR Ts have the advantages of *in vivo* expansion and accumulation, preservation of CAR expression on the T cell surface, less exhaustion, maintenance of metabolic fitness, and effector function, leading to stronger antitumor efficacy.

## Discussion

Unlike conventional anticancer drugs that kill or inhibit tumor cells unidirectionally, the interactions of the “living medicines” CAR T cells with tumor cells are reciprocal. As CAR Ts bind and kill tumor cells, they receive feedback that reshapes their own life and function. Thus, the antitumor efficacy is not determined solely by CAR T’s cytotoxicity, but also by the feedback they receive. Here, we show that engagement kinetics, especially antigen-binding off-rate, is a critical, yet underappreciated determinant of CAR T tumor infiltration and functional fitness. By decoupling *k_off_*from epitope specificity using a matched pair of EGFR-targeting mAbs, we show that fast-on/off binding kinetics has the mechanical advantages for retaining CAR molecules, infiltrating into tumor mass, and providing a necessary “kinetic escape” for CAR Ts to keep functional fitness under chronic intense antigen stimulation. While slow-off binding induces severe CAR stripping, entrapment at tumor periphery, and rapid terminal exhaustion, fast-off rate limits antigen occupancy, preserves CAR expression, drives deep tumor parenchymal infiltration, and enhances expansion and immune memory. These effects result in improved antitumor efficacy in a clinically relevant high-tumor burden model.

Previous studies, including our own^24^ and work from the Riddell lab^25^, demonstrated that low-affinity CAR T cells exhibit more tumor infiltration. However, those studies utilized distinct antibody clones and did not study the CAR T locations inside tumor mass, leaving the role of binding kinetics on intratumoral spatial distribution unresolved. To isolate these kinetic parameters, we utilized structurally paired mAbs in CAR design. While slow-off Cet CAR T cells sequestered at the tumor periphery and rapidly declined in number, fast-off H1 CAR T cells formed highly proliferative clusters deep within the tumor parenchyma. Prior models in human-antigen transgenic mice observed preferential accumulation of lower-affinity CAR T cells in tumors but attributed this to a peripheral “antigen sink” that traps high-affinity CAR T cells in healthy tissues^22^. Because our study utilized NSG mice lacking endogenous human EGFR expression, we ruled out peripheral diversion and uncovered an intrinsic, mechanical mechanism of infiltration. We propose a “pulsatile migration” model: the short dwell-time of fast-off CARs prevents physical, affinity-mediated immobilization at the tumor border. Through this rapid, iterative engagement mode, fast-off CAR T cells avoid adhesive lockdown by peripheral tumor cells or extracellular matrix, enabling efficient step-migration deep into the tumor mass.

CAR-mediated trogocytosis is increasingly recognized as a major driver of T cell dysfunction^42^. While it can occur via cell-to-cell contact without cognate binders^43^, specific high-affinity interactions drastically accelerate this membrane exchange^44^ ^45^. Trogocytosis compromises CAR T function by several mechanisms, including the acquisition of tumor antigens induces T cell fratricide^44^ and tumor-mediated receptor stripping deprives CAR T cells of signaling competence^42^, a process further exacerbated by disruptions in Rab5-dependent endocytic recycling^46^. Tumor contact indeed structurally and functionally reshapes CAR T cells^47^. On this note, the slow-off CAR T cells engage with tumor cells in a prolonged “1-on-N” mode that imposes severe physical stress, inducing ∼70% CAR downregulation that may be catastrophic to the T cells within the tumor lesion (fig. S6C vs fig. S2). In contrast, downregulation of fast-off H1 CAR is much less in tumor lesion and after *in vitro* co-culture. Importantly, fast-off H1 CAR recovers much faster than the slow off CAR. Thus, fast-off engagement limits antigen occupancy, minimizes trogocytosis, and protects CAR T cells from AICD and exhaustion. Under the translational model of treating high-burden tumors with low dose T cells, the slow-off Cet CAR Ts collapsed after transient expansion, whereas the fast-off H1 CAR Ts exhibited a highly durable, multiphasic expansion profile, surviving early pressure to mediate sustained tumor regression. This agrees with emerging data showing that moderate affinity/fast-off CAR Ts may have therapeutic benefits in addition to reducing off-tumor toxicity^6^ ^23^ ^48^.

The impact of antigen-binding affinity and kinetics in blood cancers remain controversial. While fast-off CAT-scFv (*K_D_*=14nM, *k_off_*=3.1×10^-3^s^-1^, *T_1/2_*=223s) CD19 CAR T demonstrated durable and stronger responses in leukemia patients^23^ ^48^, most of the FDA-approved CD19 targeting CARs utilized high affinity/slow-off FMC63 scFv (*K_D_*=5.1nM, *k_off_* =5.2×10^-4^s^-1^(*T_1/2_*=1332s)^49^. Furthermore, preclinical reports continue to highlight high-affinity binders with prolonged dwell times (*T_1/2_*≈3000s) that enhance CD19 CAR efficacy^50^. This discrepancy suggests that dissociation kinetics are less restrictive in liquid tumors, where a “1-on-1” engagement mode prevails. In suspension, the rapid lysis of a target cell automatically disengages the CAR T cell, granting a functional rest period that renders the intrinsic off-rate less critical.

Using single-cell transcriptomics, we further revealed that fast-off kinetics fundamentally reshape the long-term differentiation and metabolic programming of CAR Ts *in vivo*. While both CAR constructs appeared very similar following acute *in vitro* stimulation and even at early timepoints *in vivo* (Day14), a dramatic divergence emerged by Day18-20. Cet CAR Ts progressed to terminal exhaustion and shifted toward chronic aerobic glycolysis, a metabolic hallmark of persistent antigen overstimulation. Conversely, H1 CAR Ts resisted this developmental dead-end, successfully maintaining a robust pool of cytotoxic effector memory T cells sustained by OXPHOS. This metabolic plasticity, afforded by the mechanical “kinetic escape” of the H1 binder, explains the late-stage expansion rebound, and durable persistence observed exclusively in the H1 cohort.

Although our study relied on a limited number of antibody pairs due to current technological challenges of precisely tuning dissociation rates, our core conclusions are strongly supported by a fast-off F292A LFA CAR T^51^ that similarly demonstrated enhanced tumor control^52^. Beyond expanding validation across additional targets, future engineering pipelines must incorporate diverse kinetic variants to map optimal on/off thresholds. Our work establishes a conceptual framework for solid tumor CAR design. Rather than screening libraries solely for high affinity, discovery pipelines should prioritize optimal dissociation constants. This design principle builds a passive defense mechanism into the CAR, enhancing parenchymal infiltration and shielding CAR T cells from the hostile, antigen-dense microenvironment, ultimately bringing artificial CAR architecture closer to the elegant, durable design of natural TCR-mediated immunity.

EGFR is overexpressed across a broad spectrum of human malignancies^53^. Several high and low affinity CAR Ts are being developed for EGFR+^54^ ^55^ and EGFRvIII variant^56^ ^57^, but yield limited clinical responses^58–60^. Even strategies utilizing co-expressed high-affinity T-cell engagers produced only transient responses^61^. Notably, the only EGFR-targeting CAR T cells to achieve a significant clinical response^62^ have a *k_off_* of 1.12×10^-3^s^-1^ (*T_1/2_*=618s). Given that Cetuximab is already an established therapy for HNSCC and colorectal cancers, the superior preclinical efficacy, robust metabolic fitness, and projected low off-tumor toxicity of the fast-off H1 CAR T cell present a highly promising candidate for clinical translation.

## Supporting information

Suppl tables and figure for "fast-off binding...

## Acknowledgement

We dedicate this work to the memory of Ms. Yibing Peng, who initiated this project and made pivotal preliminary findings that inspired us to continue investigating the role of antigen-binding kinetics in CAR design.

We acknowledge the tremendous help of Georgia Cancer Center (GCC) Flow Cytometry Core (RRID: SCR_025747). Many of the animal studies were also assisted by GCC Small Animal Imaging Core (RRID:SCR_027047). Our studies were also helped by the expertise from Histology Core (RRID: SCR_026810) in conducting immunohistochemical staining and Integrated Genomic Core (RRID: SCR_026483) in performing the scRNAseq.

The work is partly supported by the GCC intramural Paceline Award and grant NCI R01 CA235159 to YH, grant NIH R01CA264983 to HS, and funds from Augusta University to NRJG. All funding agencies have no involvement and influence in the experimental design, data collection and analysis, and this manuscript writing and submission.

## Author contributions

YH and RM postulated the hypothesis, planned and executed the experiments, analyzed and summarized data, wrote the first draft of the manuscript that is critiqued and enhanced with all co-authors. MH conducted some of the initial plasmid construction experiments; OK and RS helped with experimental design and data analysis with their statistics expertise; HS and BK conducted the scRNAseq data analysis; NRJG and VK helped with data analysis and discussion on the applications of fast-off CAR Ts. The order of the authors was assigned based on the relative contributions each made to the research project.

## AI uses declaration

AI tools were used for manuscript English editing (Google AI mode) and for assisting with optimization and generation of the scRNA-seq analysis code (Chatgpt). All AI-assisted manuscript texts were verified word-by-word and edited as needed by authors, who take full responsibility for the content of the published article. The AI-assisted analysis code was reviewed and verified by the authors and the results were confirmed to be reproducible.

