## Supplementary material for "Fast receptor-antigen dissociation kinetics promote CAR-T cell tumor infiltration, functional fitness, and antitumor efficacy in solid tumors": Suppl tables and figure for "fast-off binding...

**Supplemental Table 1: Biophysical parameters of human EGFR-specific mAbs from literature.**

| | mAb | $k_{on}$ (1/Ms) | $k_{off}$ (1/s) | $K_D$ (nM) | $\tau_{1/2}$ (0.693/$k_{off}$) | Notes |
|---|---|---|---|---|---|---|
| Pair 1 | Cet (1) | $4.4x10^5$ | $1.42x10^{-3}$ | 3.18 | 488 | scFv-Fc format, produced in CHO |
| | H1(1) | $7.18x10^5$ | $9.27x10^{-3}$ | 13.10 | 75 | |
| Pair 2 | H528 (2) | $2.5x10^5$ | $6.0x10^{-3}$ | 25 | 115 | Fv (variable region), produced in E. coli |
| | 2HH11 (2) | $2.8x10^5$ | $2.4x10^{-4}$ | 0.86 | 2887 | |

**Supplemental Table 2: Variations of the biophysical parameters of Cetuximab reported from different groups using different antibody formats.**

| | $k_{on}$ (1/Ms) | $k_{off}$ (1/s) | $K_D$ (nM) | $\tau_{1/2}$ (0.693/$k_{off}$) | Notes |
|---|---|---|---|---|---|
| Patel et al (3) | $2.2 \times 10^5$ | $1.1 \times 10^{-3}$ | 5.2 | 630 | Original Cetuximab, produced in Sp2/0 cells |
| Reilly et al (4) | $4.9 \times 10^5$ | $1.2 \times 10^{-3}$ | 2.5 | 577 | |
| Talavera et al (5) | $3.1 \times 10^6$ | $5.8 \times 10^{-3}$ | 1.8 | 119 | Fab format (Cetuximab treated with papain) |
| Goulet et al (1) | $4.4 \times 10^5$ | $1.42 \times 10^{-3}$ | 3.18 | 488 | scFv-monoFc format produced in CHO cells |
| Zhuang et al (6) | $1.68 \times 10^5$ | $7.29 \times 10^{-5}$ | 0.4 | 9506 | Full antibody (IgG1/C*k*), produced in 293 cells |

The affinity and binding kinetics of Cetuximab to human EGFR have been previously reported in multiple studies; however, substantial variability exists across publications, likely due to differences in antibody formats and experimental conditions. Among four reports (1, 3-5), while the reported equilibrium dissociation constants ($K_D$) are broadly comparable, the dissociation rates ($k_{off}$) vary considerably. Notably, a more recent study (6) employing an scFv-Fc format reported kinetic parameters that differ markedly from earlier publications ($K_D$=0.4nM, koff = 7.29 × $E^{-5}$), further underscoring the inconsistency in reported binding kinetics.

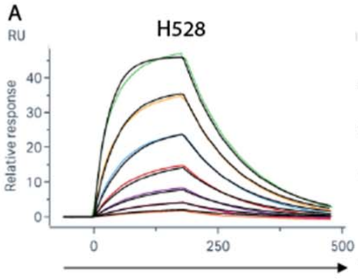

B

| Analyte | (scFv-Fc) | $k_{on}$ (1/Ms) | $k_{off}$ (1/s) | $K_D$ (nM) | $T_{1/2}$ (s) | Capture level (RU) | Relative binding response | Rmax (RU) |
|---|---|---|---|---|---|---|---|---|
| Human | H528 | $4.89 \times 10^4$ | $8.98 \times 10^{-3}$ | 184 | 77 | 73.6 | 63.41% | 63.0 |
| EGFR | 2HH11 | $4.90 \times 10^4$ | $4.56 \times 10^{-3}$ | 93.1 | 153 | 47.4 | 66.40% | 42.5 |

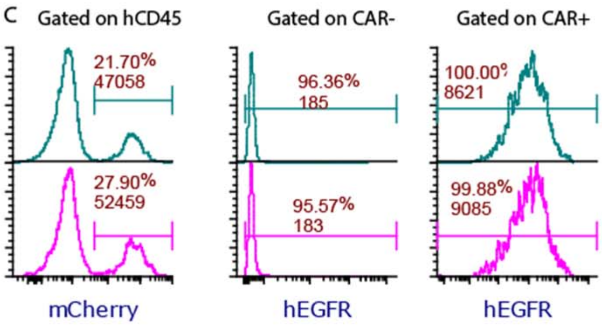

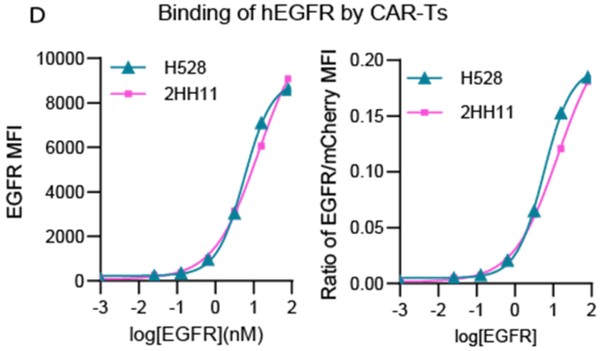

**fig. S1. Comparable biophysical and cellular binding profiles of paired H528 and 2HH11 scFv-Fc antibodies and derived CAR T cells, related to Figure 1. (A)** SPR sensorgrams showing the binding kinetics of soluble human EGFR (sEGFR) to captured H528 and 2HH11 scFv-Fc antibodies. Black lines represent global kinetic fits using a 1:1 Langmuir binding model. **(B)** Summary table of biophysical parameters derived from SPR analysis, including association rate constant ($k_{on}$), dissociation rate constant ($k_{off}$), equilibrium dissociation constant ($K_D$), dissociation half-life ($T_{1/2}$), capture level, relative binding response, and maximum binding capacity ($R_{max}$). **(C)** Representative flow cytometry histograms demonstrating comparable CAR levels (mCherry$^+$, gated on hCD45$^+$ human primary T cells) and subsequent soluble hEGFR binding profiles on gated CAR$^-$ and CAR$^+$ T cell populations for H528 (teal, top) and 2HH11 (magenta, bottom) CAR T cells. **(D)** Dose-response binding curves of soluble hEGFR to H528- and 2HH11- CAR T cells, displaying the raw EGFR MFI (left) and the normalized EGFR/mCherry MFI ratio (right) across a range of log[sEGFR] concentrations.

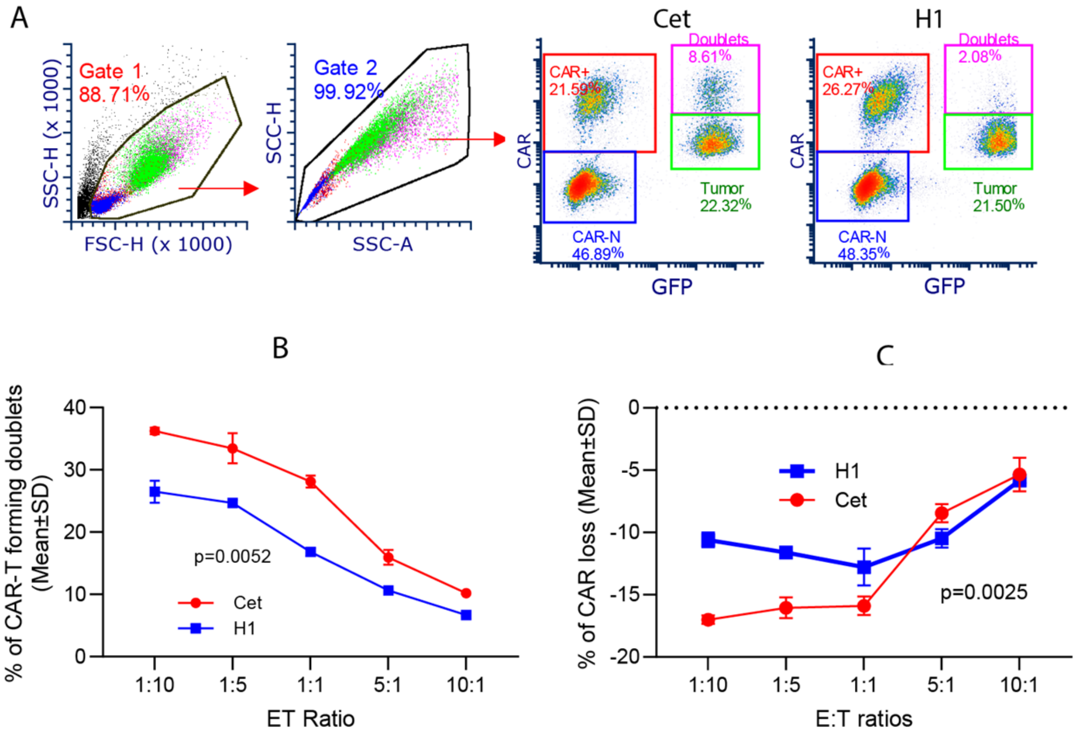

**fig. S2. Fast-off H1 CAR T cells exhibit reduced doublet formation with tumor cells and reduced CAR loss *in vitro*, related to Figure 2.** The experimental setup was identical to that described in Figure 2, utilizing primary human CAR T cells in place of Jurkat cells. **(A)** Flow cytometry gating cascade and representative dot plots identifying cell doublets (magenta gate) formed between Cet or H1 CAR T cells and GFP$^+$ A431 tumor cells. Sub-gates delineate CAR$^+$ T cells (red), CAR$^-$ T cells (blue), and unengaged tumor cells (green). **(B)** Quantification of the percentage of CAR T cells participating in doublet formation across the indicated effector-to-target (E:T) ratios following a 30-min incubation at 37°C. **(C)** Downregulation of CAR molecules on CAR T cells, expressed as the percentage loss of mCherry MFI following co-culture with target tumor cells. Data are presented as mean ± SD from triplicates. Statistical significance in **(B)** was determined by two-way ANOVA (p-value indicated on plot).

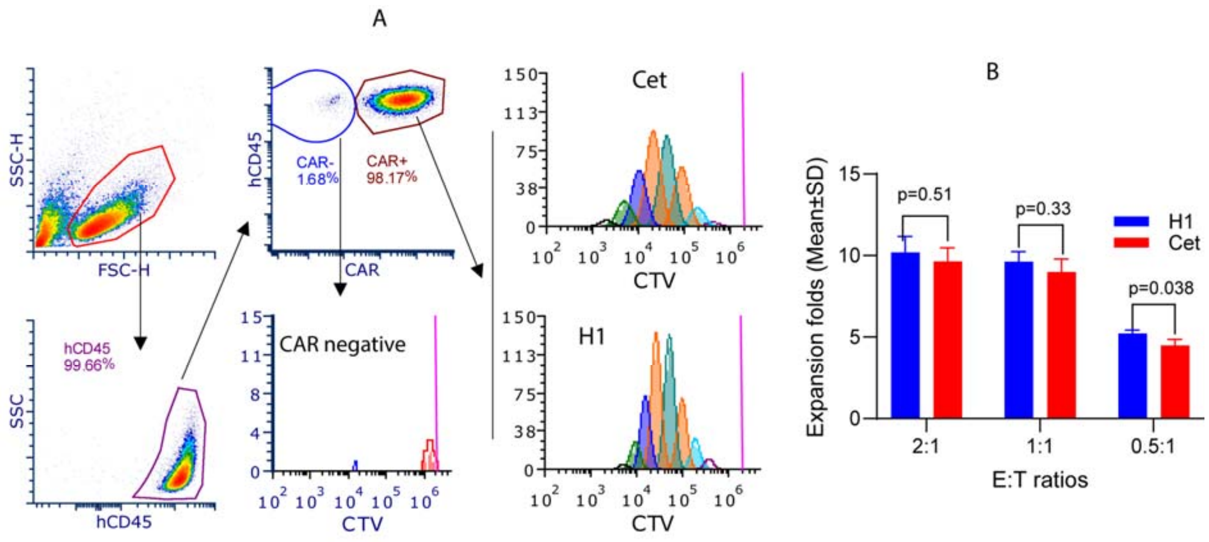

**fig. S3. Slow-off Cet and fast-off H1 CAR T cells exhibit comparable proliferation dynamics upon co-culture with EGFR$^{moderate}$ FaDu tumor cells, related to Figure 3. (A)** Flow cytometry gating strategy and representative histograms tracking CAR T cell proliferation via CellTrace Violet (CTV) dye dilution following co-culture with FaDu cells. Cells were sequentially gated on live cells, hCD45$^{+}$ human T cells, and subsequently segregated into CAR$^{-}$ (blue gate) and CAR$^{+}$ (orange gate) populations. Proliferation profiles are displayed for the unstimulated CAR$^{-}$ control population (bottom center) and antigen-engaged Cet (top right) or H1 (bottom right) CAR$^{+}$ T cells. Successive multi-colored peaks represent distinct cellular generations resulting from division, with the undivided baseline indicated by the magenta vertical line. (B) The expansion folds were calculated by dividing the total CAR T cells after coculture with tumor cells with the initial input number of CAR T cells. Data are representative of three independent experiments.

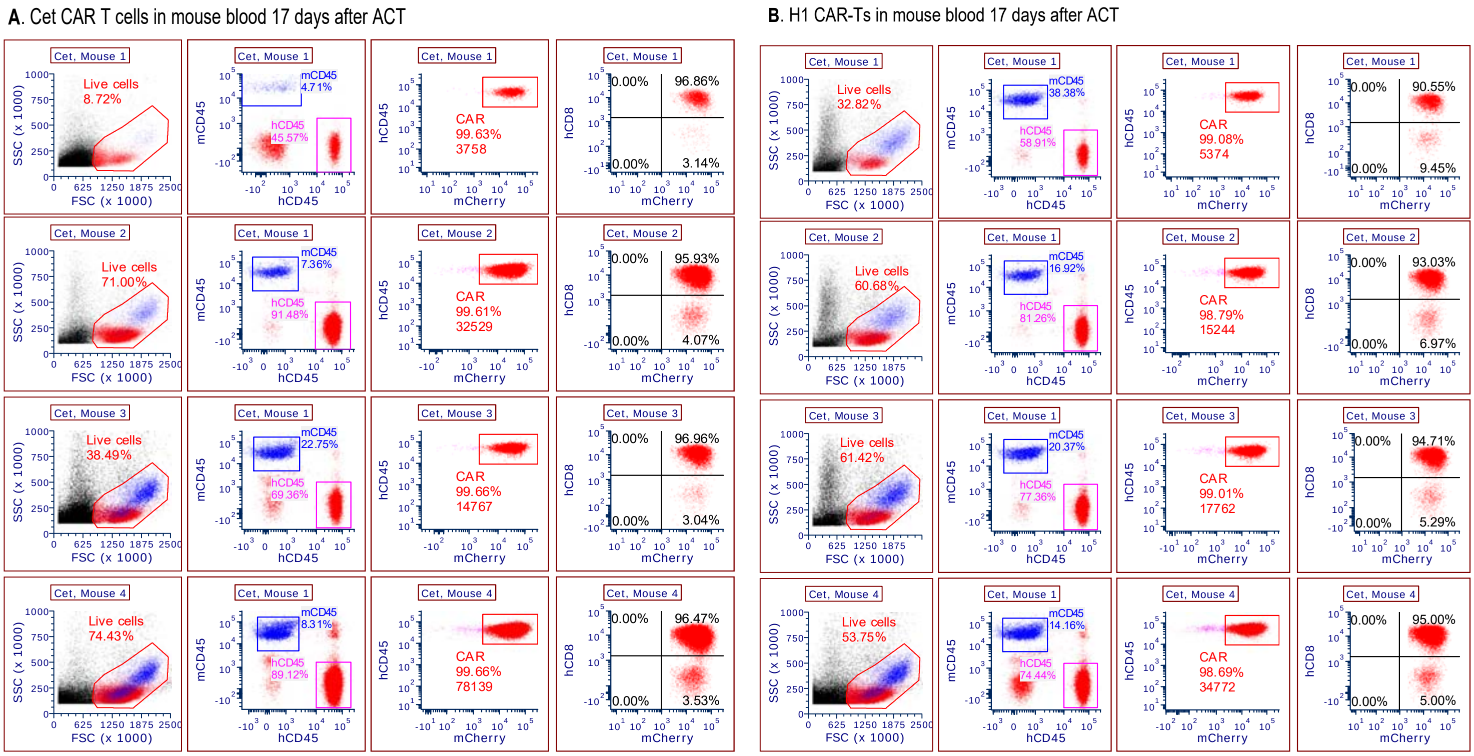

**fig. S4. Flow cytometry data of tracking CAR T expansion in individual mouse blood, related to Figure 4.** Flow cytometry analysis of peripheral blood from individual mice (Mice 1–4) on day 17 post-adoptive cell transfer to quantify human CAR T cells. **(A and B)** Gating cascades are shown for mice treated with Cet CAR T cells **(A)** or H1 CAR T cells **(B)**. For each individual mouse, peripheral blood mononuclear cells were sequentially analyzed through a multi-step gating strategy: **First Panel (Left):** Identification of the total viable leukocyte population based on forward scatter (FSC) and side scatter (SSC) characteristics. **Second Panel:** Relative of human and mouse leukocytes by gating on hCD45$^{+}$ cells (magenta gate) versus mouse mCD45$^{+}$ internal control cells (blue gate). **Third Panel:** % of mCherry$^{+}$) (CAR$^{+}$) and CAR T number. **Fourth Panel (Right):** the % of CD8$^{+}$ T cells among the CAR T cell population.

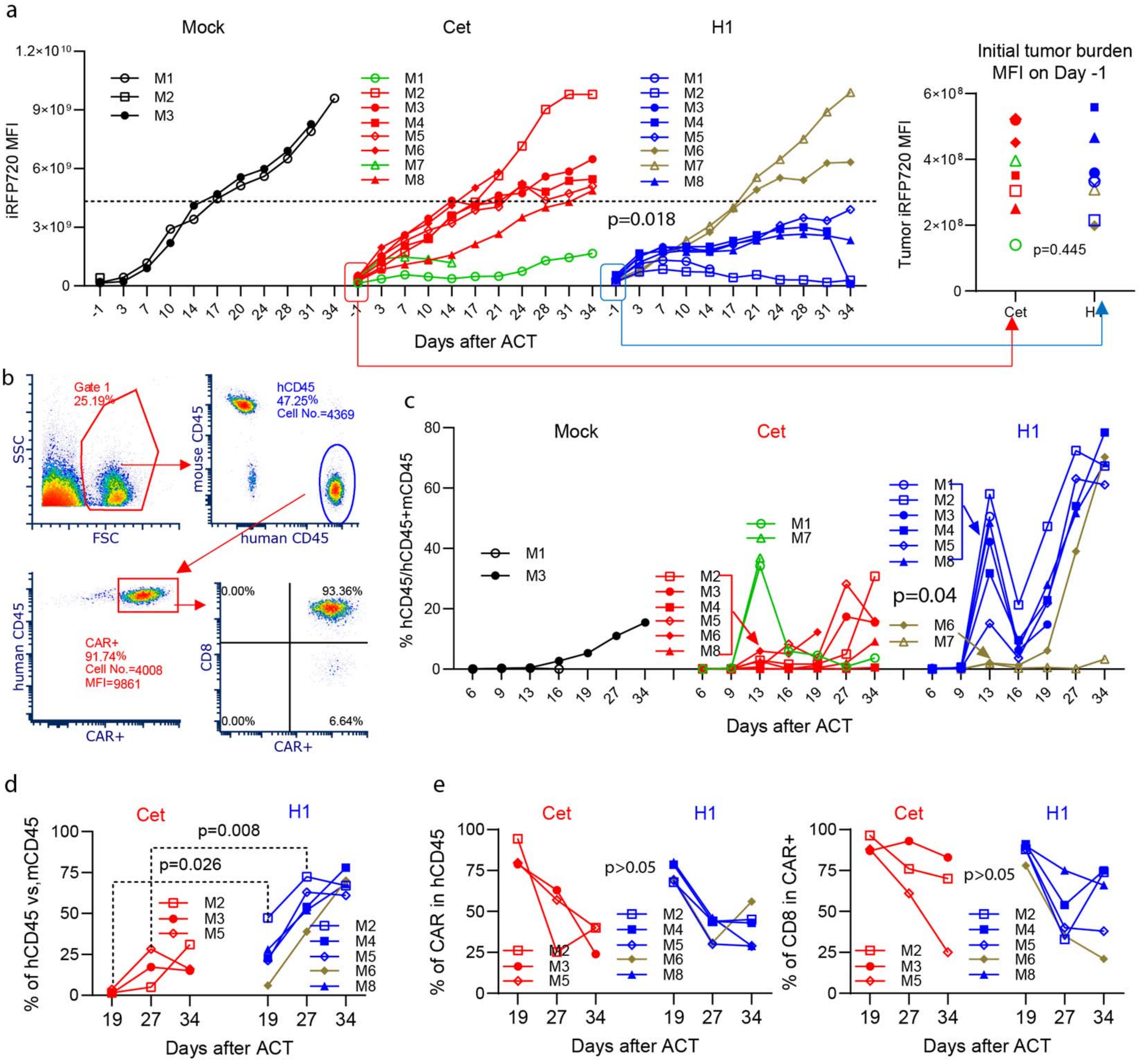

**fig. S5A. Individual mouse tumor kinetics and peripheral blood CAR T cells, related to Figure 5. (a)** Longitudinal monitoring of individual tumor burdens by iRFP720 MFI at indicated days after ACT. The horizontal dashed line defines a threshold distinguishing mice exhibiting therapeutic effects and substantial CAR T cell expansion (below the line) from non-responding animals (above the line). Statistical significance was determined by a Wilcoxon signed-rank test (p=0.018). **(b)** Flow cytometry gating strategy evaluating blood CAR T cell expansion. Viable leukocytes (Gate 1) were gated to discriminate $hCD45^+$ T cells (blue oval) from mouse $CD45^+$ cells, followed by $CAR^+$ expression ($mCherry^+$, red rectangle) and % of $hCD8^+$ in the CAR+. **(c)** Kinetics for individual mice tracking blood human T cell expansion, expressed as the percentage of $hCD45^+$ cells within total human and mouse leukocytes (% $hCD45^+$/$hCD45^+$ + $mCD45^+$). Statistical analysis comparing treatment groups was performed using Welch's two-sample t-test on log-transformed area under the curve (log-AUC) values (p=0.04). Note: At day 13, significant CAR T expansion and tumor control were detected in 6/8 mice (75%, blue lines) in the H1 cohort, compared to only 2/8 mice (green lines, 25%) in the Cet group. (d) Re-expansion of CAR Ts is reflected by a second peak in hCD45+ cells. (e) The concurrent decrease in the %of CAR+ and CD8+ T cells indicates the onset of GvHD. Individual mouse symbols match across panels to correlate antitumor efficacy with CAR T expansion.

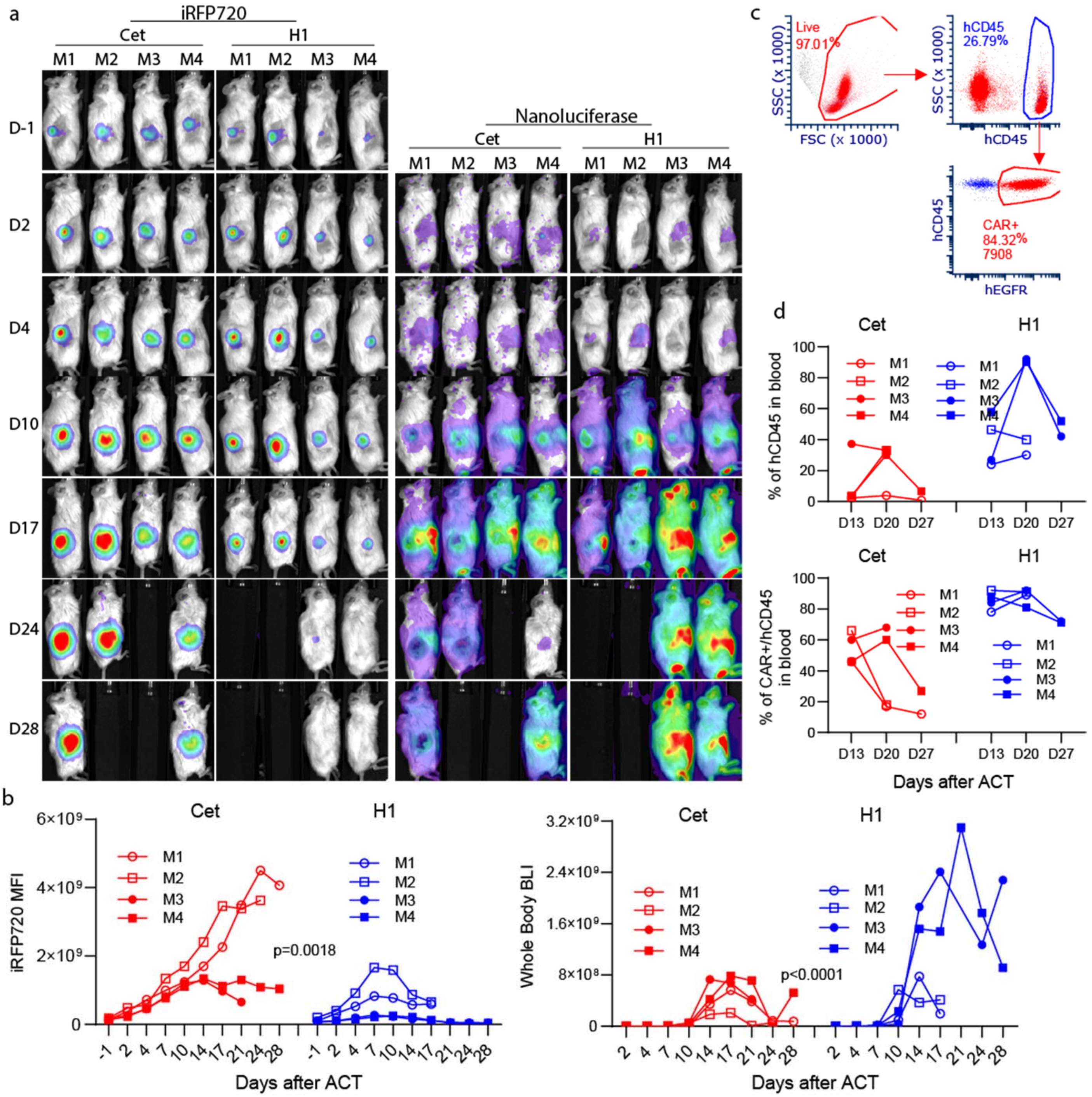

**fig. S5B. Dual-reporter imaging reveals enhanced *in vivo* expansion and antitumor efficacy of fast-off H1 CAR T cells, related to Figure 5.** NSG mice were subcutaneously inoculated with $2x10^6$ A431-iRFP720 tumor cells. Eight days post-inoculation, mice were adoptively transferred with $3x10^6$ Cet or H1 CAR T cells co-expressing NanoLuciferase (NanoLuc). Tumor progression and spatial CAR T biodistribution/expansion were longitudinally monitored via concurrent iRFP720 fluorescent and bioluminescent imaging, alongside peripheral blood flow cytometric analysis. **(a)** Serial near-infrared fluorescent whole-body images mapping tumor burden (iRFP720, left) alongside corresponding BLI tracking systemic CAR T cell distribution and expansion kinetics (NanoLuciferase activity, right) in individual mice (M1–M4) across treatment cohorts. **(b)** Kinetic trajectories for individual mice quantifying longitudinal tumor growth (iRFP720 MFI, left) and whole-body CAR T cell expansion (Whole-Body BLI total flux, right). Statistical significance between cohorts over time was determined by two-way ANOVA ($p=0.0018$ for tumor burden; $p<0.0001$ for CAR T expansion). **(c)** Representative flow cytometry gating cascade tracking circulating human T cell. Viable leukocytes (Live gate) were interrogated to identify $hCD45^+$ T cells (blue gate), followed by downstream resolution of the $mCherry^+$ CAR T cell fraction (red gate). **(d)** Individual single-mouse kinetic tracking of peripheral blood human T cells, displayed as the percentage of $hCD45^+$ leukocytes out of total leukocytes (top graph), and % CAR, displayed as the percentage of $mCherry^+$ cells within the hCD45+ T cell fraction (bottom graph) on days 13, 20, and 27 post-ACT.

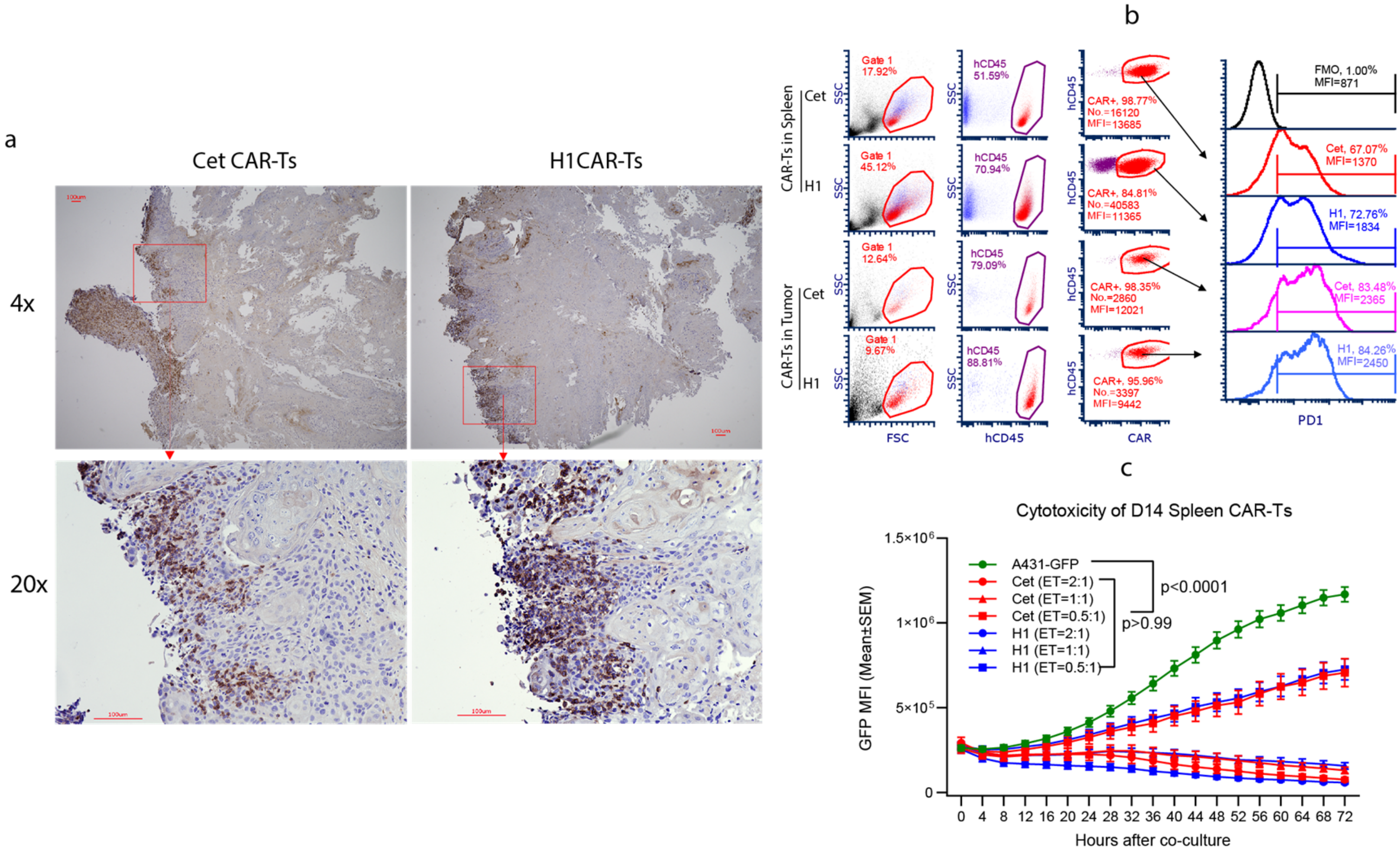

**fig. S6A. Spleen- and tumor-derived Cet and H1 CAR T cells maintain comparable functional capacity at an early *in vivo* checkpoint (Day 14 post-ACT). (a)** IHC staining of the tumor tissues collected at D14 after treatment with either Cet or H1 CAR T cells. **(b)** Flow cytometric histogram profiles tracking PD-1 expression levels on Cet and H1 CAR T cells isolated from the spleen (top panels) and tumor lesions (bottom panels) on day 14 (D14) after ACT. Fluorescence minus one (FMO) gating controls are indicated in black at the top of each stack. **(c)** *Ex vivo* real-time cytotoxicity kinetics assessing the tumor-killing capacity of D14 spleen-derived (left) and tumor-infiltrating (right) Cet and H1 CAR T cells against $GFP^{+}$ A431 target cells by IncuCyte real time cell analyzer. Data are representative of two independent experiments and are presented as mean ± SEM. Statistical significance for killing kinetics was determined by two-way ANOVA (p-values are indicated within the panels).

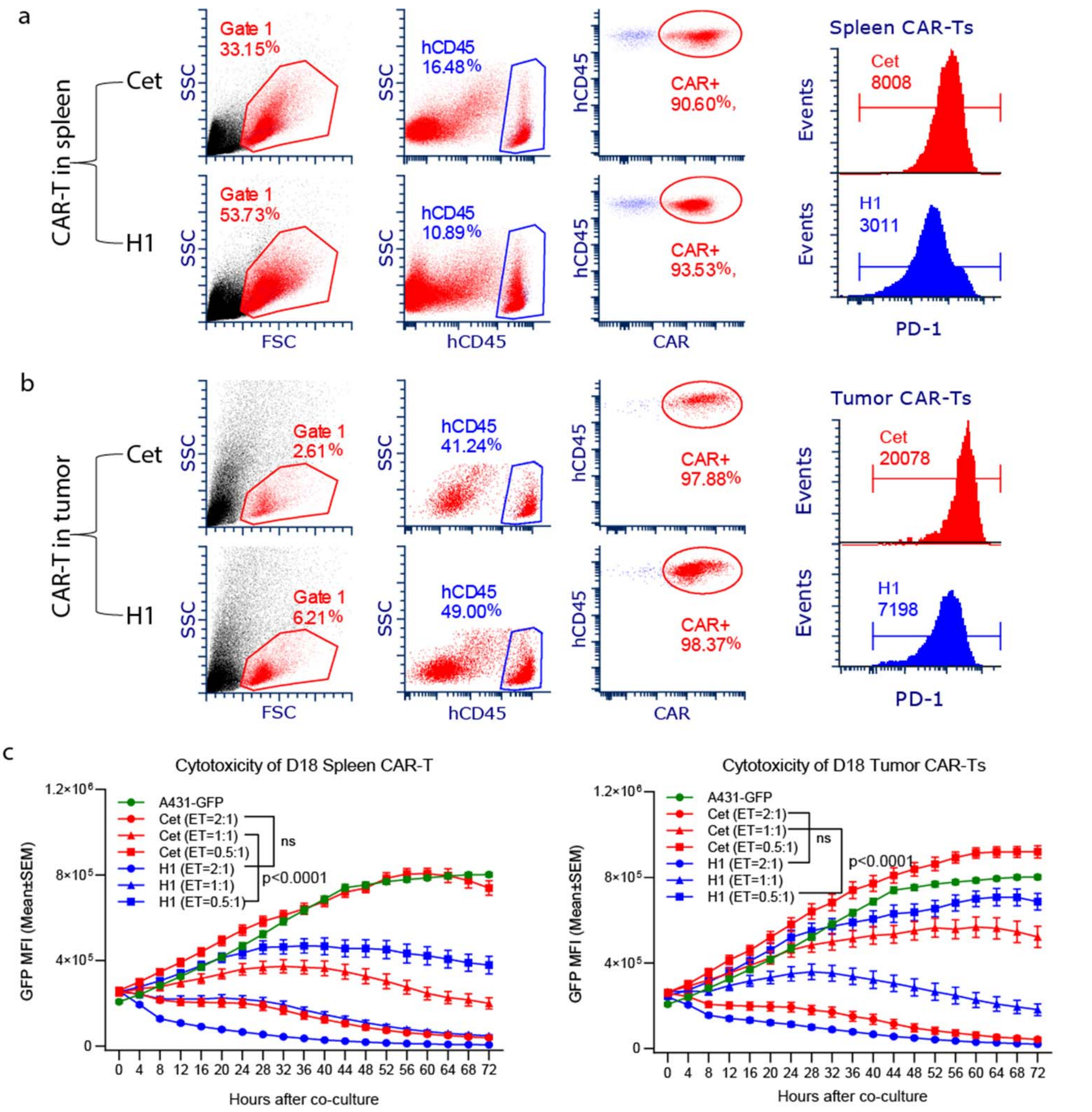

**fig. S6B. Fast-off H1 CAR T cells exhibit reduced exhaustion and superior functional durability at an intermediate *in vivo* checkpoint (Day 18 post-ACT), related to Figure 6.** Spleen tissues and tumor lesions were harvested on day 18 (D18) after ACT to evaluate CAR T cell phenotyping and functional preservation. **(a and b)** Representative flow cytometry gating strategies and histogram profiles evaluating PD-1expression levels on Cet and H1 CAR T cells isolated from the spleen **(a)** and tumor mass **(b)**. For both tissue compartments, viable leukocytes (Gate 1, red gate) were tracked to identify hCD45$^+$ cells (blue gate), followed by downstream isolation of the CAR T cell fraction (red oval). Right histograms display relative PD-1 expression frequency and MFI values (Cet in red; H1 in blue). **(c)** *Ex vivo* real-time cytotoxicity kinetics assessing the tumor-killing capacity of D18 spleen-derived (left) and tumor-infiltrating (right) Cet and H1 CAR T cells against GFP+ A431 target cells at the indicated E:T ratios over 72 h using an IncuCyte live-cell imager. Data are representative of two independent experiments and are presented as mean ± SEM. Statistical significance for killing kinetics was determined by two-way ANOVA (p-values and non-significance [ns] markers are indicated within the panels).

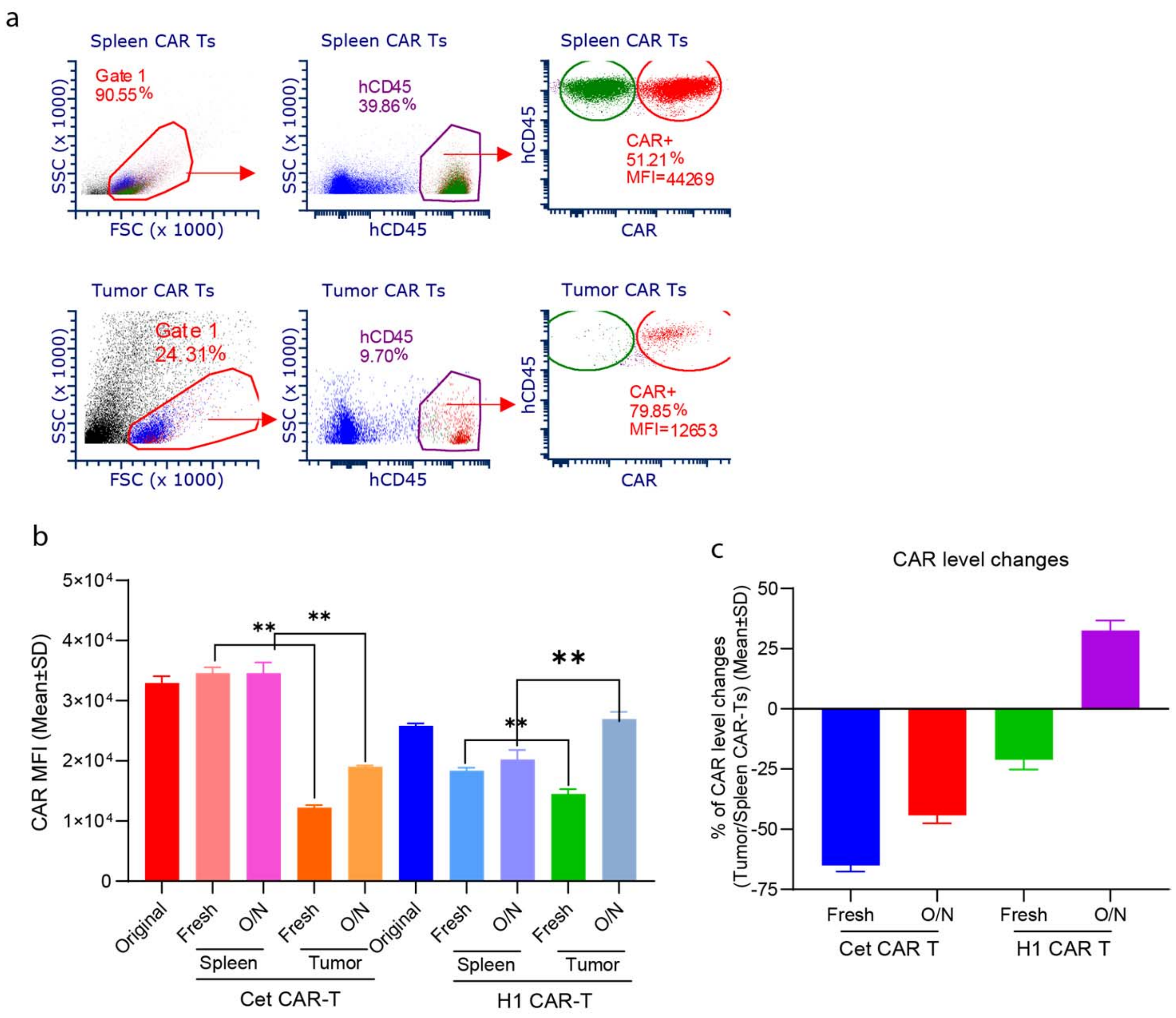

**fig. S6C. Fast-off H1 CAR T cells resist antigen-induced CAR downregulation within the tumor microenvironment and exhibit rapid receptor recovery. (a)** The gating strategy to analyze the CAR levels of the CAR Ts from spleen and tumor. (**b**) Flow cytometric quantification of CAR levels (MFI) on Cet and H1 CAR T cells. CAR surface expression was analyzed either immediately upon isolation ("Fresh") or following overnight culture ("O/N") in media supplemented with low-dose interleukin-2 (IL-2) and compared back to baseline un-transferred parental cells ("Original") across spleen and tumor tissue compartments. **(c)** Percentage change in CAR surface levels on tumor-infiltrating lymphocytes (TILs) relative to their corresponding splenic CAR T cell counterparts under fresh or overnight-rested ("O/N") culture conditions. Data are presented as mean ± SD from triplicates. Statistical significance was determined by an unpaired Student's t-test (** indicates $p < 0.001$).

A

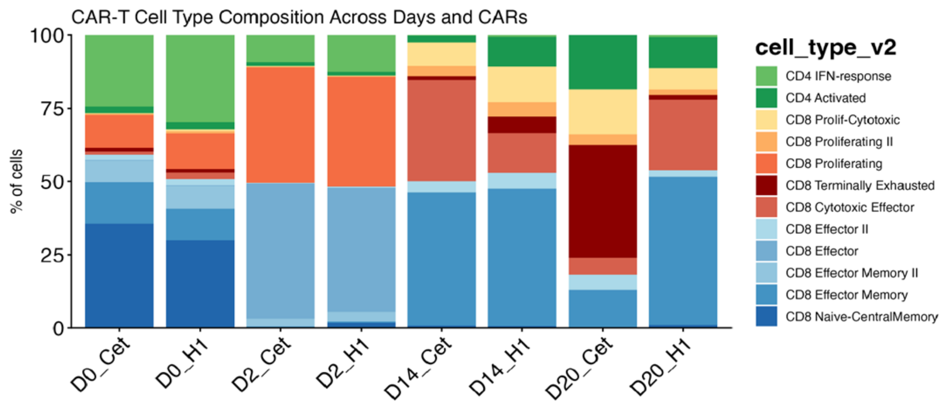

B

| Group | CD8 Terminally Differentiated | CD8 Effector Memory |
|---|---|---|
| Cet_D20 | **~75%** | ~10% |
| H1_D20 | ~5% | **~45%** |
| Cet_D14 | ~5% | ~50% |
| H1_D14 | ~12% | ~45% |

C

| Program | H1 higher (negative LFC) | Cet higher (positive LFC) |
|---|---|---|
| Memory/Stemness | CCR7 (−2.82), IL7R (−2.27), TCF7 (−1.85), LEF1 (−1.91), IL2 (−2.71), BACH2, LRRN3 | — |
| Exhaustion/Terminal | — | HAVCR2 (+2.41), TOX2 (+3.01), EOMES (+3.57), LAG3 (+1.35), PDCD1 (+1.18), KLRG1 (+1.82) |
| Effector | — | GZMK (+1.93), CX3CR1 (+2.23), IFNG (+1.94), PRF1 (+1.25) |
| TF program | FOXO1, KLF2, S1PR1 | TBX21, PRDM1 |

D

| pathway | D14 | D20 |
|---|---|---|
| Glycolysis (18 genes) | **H1 higher** (PFKFB4 −3.26, ALDOC −2.36) | **Cet higher** (PFKFB4 +1.85, ALDOC +1.61) |
| Hypoxia/HIF1A | **H1 higher** (DDIT4 −3.42, BNIP3 −2.36, VEGFA −2.45) | **Cet higher** (DDIT4 +2.03, BNIP3 +1.52, EPAS1 +1.57) |
| OXPHOS (13 genes) | **Cet higher** (NDUFS6 +1.39) | **H1 higher** (NDUFS6 −1.27) |
| mTOR | Cet higher | **H1 higher** (MTOR −1.03) |

**fig. S7. Phenotypic and transcriptomic dynamics of fast-off H1 and slow-off Cet CAR T cells *in vitro* and *in vivo*, related to Figure 7. (A)** Bar plot illustrating the longitudinal CAR T cell type composition and relative percentages of defined $CD4^+$ and $CD8^+$ T cell populations across time points and CAR constructs (Cet vs. H1). D0: baseline CAR T cells prior to target cell stimulation; D2: CAR T cells following 2 days of *in vitro* tumor cell co-culture; D14 and D20: splenic CAR T cells isolated on days 14 and 20 post-adoptive transfer. **(B)** Summary table quantifying the approximate percentages of $CD8^+$ Terminally Differentiated (Exhausted) and $CD8^+$ Effector Memory T cell populations in Cet and H1 groups at D14 and D20 *in vivo*. **(C)** Summary table highlighting key differentially expressed genes (DEGs) and transcription factor (TF) programs involved in memory/stemness, exhaustion/terminal differentiation, and effector functions. $Log_2$ fold change (LFC) values indicate genes significantly enriched in H1 (negative LFC, left) or Cet (positive LFC, right) CAR T cells. **(D)** Comparative pathway summary table detailing the regulation of metabolic and signaling cascades (Glycolysis, Hypoxia/HIF1A, OXPHOS, and mTOR) between H1 and Cet CAR T cells at early (D14) and late (D20) *in vivo* time points, noting representative differentially regulated genes and their respective LFC values.
